# A proteome-wide dependency map of protein interaction motifs

**DOI:** 10.1101/2024.09.11.612445

**Authors:** Sara M. Ambjørn, Bob Meeusen, Johanna Kliche, Juanjuan Wang, Dimitriya H. Garvanska, Thomas Kruse, Blanca Lopez Mendez, Matthias Mann, Niels Mailand, Emil P.T. Hertz, Norman E. Davey, Jakob Nilsson

**Affiliations:** Center for Epigenetic Cell Memory, Danish Cancer Institute, Copenhagen, Denmark; Novo Nordisk Foundation Center for Protein Research, ICMM, University of Copenhagen, Copenhagen, Denmark; Division of Cancer Biology, The Institute of Cancer Research, 237 Fulham Road, SW3 6JB, London, UK; Department of Clinical Biochemistry, Copenhagen University Hospital Bispebjerg and Frederiksberg Hospital, Copenhagen, Denmark; Skape Bio ApS, Copenhagen, Denmark; Department of Cell and Molecular Biology, St. Jude Children’s Research Hospital, 262 Danny Thomas Place, Memphis, TN 38105, US

## Abstract

Short linear motifs (SLiMs) are the most ubiquitous protein interaction modules within unstructured regions of the human proteome, yet their contribution to cellular homeostasis remains poorly understood. To systematically assess SLiM function, we applied base editing to mutate all reported and a set of computationally predicted SLiMs defined by SLiM-like evolutionary patterns. By screening 7,293 SLiM-containing regions with 80,473 mutations in HAP1 cells, we define a SLiM dependency map identifying 450 known and 264 predicted SLiMs required for normal cell proliferation. Mutational consequences were highly reproducible in RPE1 cells, with differences attributed to cell line-specific gene essentiality. We show that most essential predicted SLiMs belong to novel classes and identify binding partners for several of these, providing mechanistic insight into a disease-associated ANKRD17 mutation. Our study provides a proteome-wide resource on SLiM essentiality and uncovers numerous uncharacterized yet essential SLiMs.

## Introduction

The molecular details of how specific protein complexes contribute to cellular functions is a fundamental question in cell biology and forms the basis for interpreting disease causing mutations. Short linear motifs (SLiMs; also referred to as molecular recognition features (MoRFs), molecular recognition elements (MREs), or eukaryotic linear motifs (ELMs)) are an important and undercharacterized class of protein interaction modules, likely constituting the largest uncharted part of the human interactome^1–3^. SLiMs reside in intrinsically disordered regions (IDRs), protein segments that do not adopt a stable three-dimensional structure, though they often form stable secondary structure in their bound state. They are typically encoded in short stretches of ∼10 amino acids where 2-4 residues serve as key binding determinants, directly interacting with binding partners, which are most commonly globular protein domains (Fig. 1a-b), but can also include DNA, RNA, and lipid membranes^4–6^. Due to the compact footprint of these interfaces, SLiMs generally bind with low micromolar affinity and mediate dynamic, conditional, yet specific protein interactions^7^. The simple binding determinants of SLiMs also facilitate the acquisition of motifs by *ex nihilo* evolution through random mutation, resulting in multiple proteins convergently evolving a similar SLiM that binds the same motif-binding partner^8^. Likewise, SLiM function can readily be perturbed by single amino acid mutations and this can contribute to human diseases^2,9,10^. The unique properties of SLiM-mediated interactions are widely utilized by the cell, and SLiMs play a wide range of roles to support key cellular functions^2^, including driving the assembly of dynamic complexes, recruiting modifying enzymes, controlling protein stability, and encoding subcellular localisation^5^.

**Figure 1.**
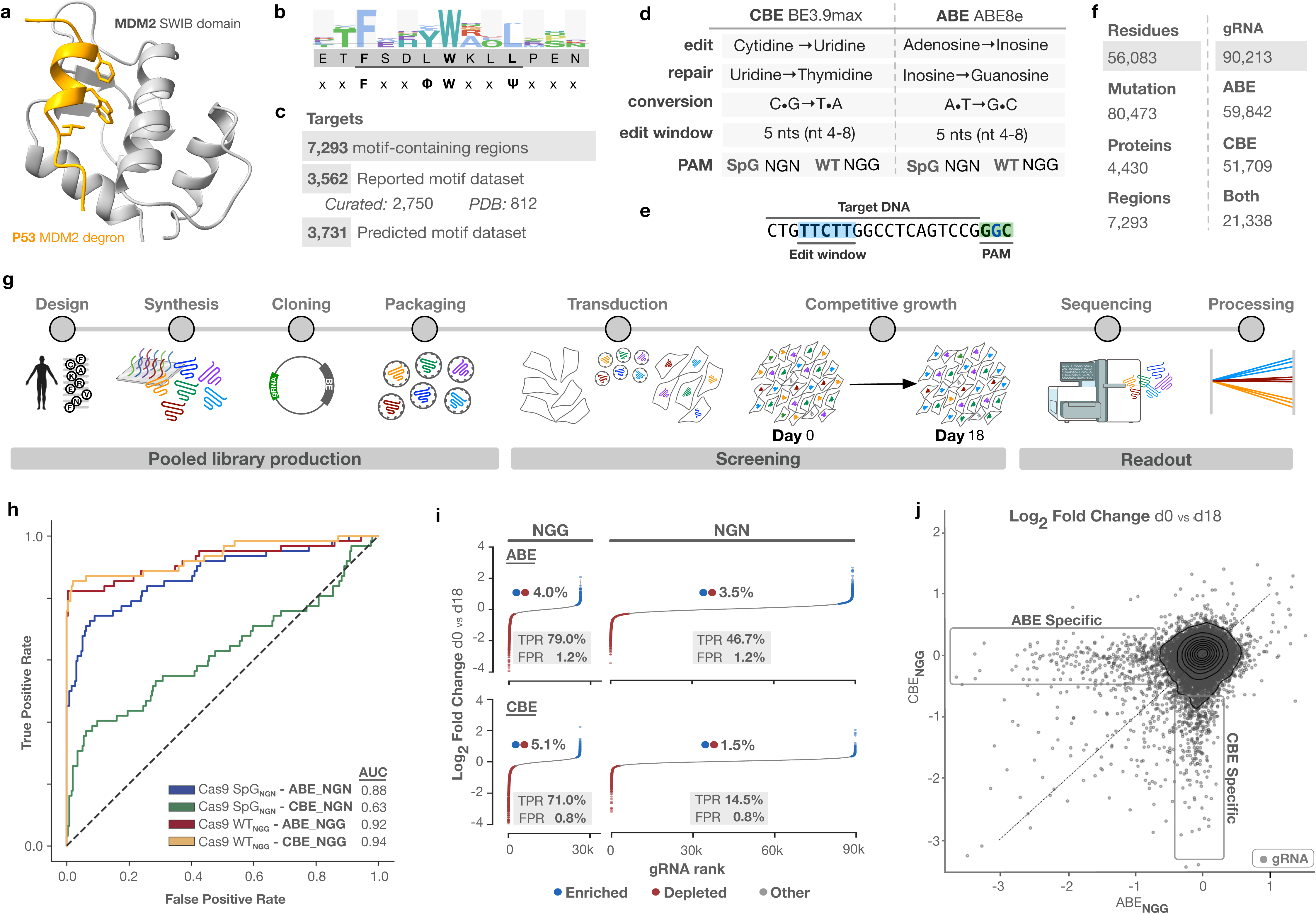
Design and benchmarking of a SLiM-targeting base editor screen. **a**, SLiM-mediated interaction exemplified by the p53 MDM2 degron binding to the MDM2 SWIB domain. **b**, Sequence logo (top), peptide sequence (middle) and consensus (bottom) of the MDM2 degron in p53, showing the evolutionary conservation of key affinity and specificity determintants (defined positions) and non-conserved flanking residues (wildcard positions denoted with ‘x’). **c**, Composition of the SLiM-targeting base editor library. **d**, Editing properties of the Cytosine Base Editor (CBE) and Adenine Base Editor (ABE) classes used in the screen. **e**, Example of a gRNA target DNA sequence with optimal base editing window and protospacer adjacent motif (PAM) highlighted. **f**, Statistics of predicted mutations introduced in target SLiMs by gRNAs cloned into ABE and/or CBE. **g**, Schematic of the base editing screening experimental pipeline. **h**, Receiver Operating Characteristic (ROC) curve based on the gRNA fold change for the positive and negative control gRNAs across the four base editors. AUC, area under the curve. **i**, Rank plots of the gRNA log_2_ fold change showing the significantly enriched, depleted, and unchanged gRNAs. TPR: True positive rate. FPR: False positive rate. **j**, Comparison of the gRNA log_2_ fold change in the ABE_NGG_ and CBE_NGG_ screens.

Decades of detailed research has uncovered thousands of human SLiMs, with the potential for several thousand more to be discovered^1^. The recent development and application of a range of novel high-throughput methods to identify peptides binding to a globular domain has increased the rate of SLiM discovery^3^. Although detailed mechanistic and functional studies have been conducted for a small subset of the known SLiMs, most have not been functionally characterized. Functional analysis remains a time-consuming bottleneck, and a significant portion of SLiMs are validated solely by *in vitro* experiments, biophysical measurements or structural studies. Functional characterization experiments often solely rely on exogenous expression of proteins, which does not address function at the endogenous protein level. Since the affinities of SLiMs are finely tuned for proper biological function, it is crucial that SLiMs are disrupted in the endogenous proteins to maintain physiological concentrations and spatiotemporal expression^7,11^. Thus, most SLiMs are incompletely characterized, leaving a gap in our understanding of SLiM contributions to cell fitness. Approaches to functionally characterize SLiMs at scale would help in the interpretation of the large amount of interaction and disease mutation data, and pinpoint key interactions required for cellular functions. Here, we utilize a base editor screening approach to perform large-scale, programmable mutagenesis of all reported and thousands of computationally predicted SLiMs in their endogenous cellular context to investigate their impact on cellular proliferation.

## Results

### Design of a base editor screen targeting SLiMs in the human proteome

As our target set, we generated a list of 7,293 SLiM-containing regions in 4,430 proteins, where each region contains one or more overlapping motifs (Fig. 1c, Table S1). This dataset contains all reported SLiMs (the *reported motif set*) and an extensive set of predicted SLiMs (the *predicted motif set*). The *reported motif set* consists of 3,562 motif-containing regions comprising a comprehensive set of 2,750 curated motifs from the experimental literature (compiled in the Eukaryotic Linear Motif resource (ELM)^5^ and from manual curation) and 812 motif instances extracted from PDB structures. The *predicted motif set* contains 3,731 protein regions with SLiM-like evolutionary patterns predicted by screening for highly conserved groups of amino acids in rapidly evolving disordered regions (see methods section).

We reasoned that base editors would be ideal to precisely disrupt SLiM-mediated interactions by introducing point mutations in key residues required for binding, allowing the functional interrogation of SLiMs at scale and at endogenous protein levels. Base editors are composed of a catalytically impaired Cas protein (typically Cas9 nickase) fused to either a cytosine or adenine deaminase. The gRNA RNA (gRNA) contains a 20 nucleotide RNA sequence which gRNAs the Cas enzyme to the complimentary location in the genome and displaces the opposite DNA strand, which is then deaminated by the deaminase. In this manner, followed by replication and repair, cytosine base editors (CBEs) convert C-G base pairs to T-A while adenine base editors (ABEs) convert A-T base pairs to G-C^12^ (Fig. 1d). Cas binding also requires the presence of a protospacer adjacent motif (PAM), which for SpCas9 is an NGG sequence located immediately downstream of the gRNA binding sequence. Editing takes place within a specific nucleotide “editing window” at a defined distance from the PAM (Fig. 1e). This allows for gRNA RNA (gRNA) directed mutations in the genome^12^, with the amino acid mutational outcome dictated by the underlying codon sequence. In this study, we selected the ABE ABE8e^13^ and the CBE BE3.9max^14^, both of which target a 5-nucleotide editing window spanning nucleotides 4-8 in the target DNA matching the gRNA (Fig. 1e). These were fused to either SpCas9 nickase (D10A) or the engineered variant SpG nickase^15,16^, which recognizes a relaxed NGN PAM sequence, allowing for an expanded range of targetable residues in SLiMs. We will henceforth refer to these base editors as: ABE_NGG_, CBE_NGG,_ ABE_NGN_, and CBE_NGN_.

A set of gRNAs was designed to target each SLiM with the four base editors. For the prediction of mutations, we assumed full editing within the editing window and retained only gRNAs predicted to result in non-synonymous mutations. The resulting gRNA library consisted of 90,213 gRNAs targeting 80,473 unique mutations to 56,083 residues in the 7,293 target SLiMs (Fig. 1f, Table S2). The library also contained 62 previously described positive control gRNAs targeting splice sites of essential genes and 255 negative controls including gRNAs targeting intergenic regions of the genome and non-targeting gRNAs that do not map to the genome^16^. Only gRNAs targeting NGG PAMs were cloned into lentiviral vectors encoding SpCas9-base editor fusions (ABE_NGG_ and CBE_NGG,_) while all gRNAs targeting NGG or NGN PAMs were cloned into lentiviral vectors encoding SpG-base editor fusions (ABE_NGN_ and CBE_NGN_) to produce four lentiviral plasmid libraries.

### Design and benchmarking of the SLiM dependency screen

HAP1, a near-haploid human cell line derived from a chronic myelogenous leukemia (CML) cell line, had shown superior performance in previous base editor screens^16,17^ and was chosen as the cell model to interrogate SLiM dependency. HAP1 cells were transduced in biological triplicates with each of the lentiviral libraries at low multiplicity of infection and high gRNA coverage (>675-fold) (Fig. 1g). After transduction and selection, cells were grown competitively for 21 days with passaging every three days. Genomic DNA was extracted at day 0, 18, and 21 post selection, and gRNA abundances were quantified by next generation sequencing (NGS) (Fig. 1g). The fold changes in gRNA abundances comparing early and late time points were quantified along with the significance of that change using the limma gene expression analysis package to identify gRNAs which were enriched, stable, or depleted over the time course of the screen^18^ (Table S3).

Positive and negative control gRNAs were examined to assess the discriminatory power of each base editor to identify functionally relevant edits. We observed substantial depletion of positive control gRNAs compared to negative control gRNAs (Fig. S1a) with good discriminatory power for three out of four base editors (Fig. 1h, ABE_NGN_ AUC=0.88; ABE_NGG_ AUC=0.92; CBE_NGN_ AUC=0.63; CBE_NGG_ AUC=0.94). The CBE_NGN_ base editor displayed a notably weaker depletion of positive controls, consistent with observations by others^16^, which was also reflected in lower discriminatory power. Base editors targeting the NGG PAM outperformed those targeting the NGN PAM (Fig. 1h). As the discriminatory power at day 18 was highly similar to that at day 21, we focused on day 18 in the following analyses (Fig. S1b). A p-value cut-off of 0.01 showed a false positive rate of ∼1% across all four screens, with a true positive rate of up to 79% (Fig. 1i), and was set as the threshold for considering gRNAs to significantly influence cell proliferation.

We next examined the effect of the SLiM-targeting gRNAs. The majority of these showed minimal to no change in abundance during the screen; however, a small proportion exhibited significant depletion or enrichment (Fig. 1i; ABE_NGN_ 3.5%, ABE_NGG_ 4.0%, CBE_NGN_ 1.5%, and CBE_NGG_ 5.1%). We observed a strong correlation between replicates as well as between the gRNAs that changed significantly in both NGN and NGG screens for the same editor (Fig. S1c-h). As anticipated, given the different targeting and editing outcomes of the CBE and ABE, we observed distinct gRNA changes for each editor, advocating the use of both for deeper functional investigation (Fig. 1j).

We next confirmed that the base editors introduced the predicted mutations by focusing on a SLiM from the *predicted motif set* in Cleavage and polyadenylation specificity factor subunit 1 (CPSF1) that was required for cell fitness. We deep sequenced the genomic locus after transduction with 15 single gRNAs in either ABE_NGN_ or CBE_NGN_ base editors (Fig. S2). The ABE_NGN_ gRNAs introduced mutations with high efficiency at day 0 (up to 61% allelic frequency) with good overlap with the predicted edits and showed limited bystander edits (Fig. S2, Table S4 and Supplemental data). The CBE_NGN_ gRNAs also introduced the predicted mutations, though at lower efficiency than the ABE_NGN_, displaying up to 10% indel generation and evident bystander editing, which is in line with findings by others^14^ (Fig. S2, Table S4 and Supplemental data). We further observed that specific mutations introduced by gRNAs that scored in the proliferation screen were depleted from the cell population from day 0 to day 18, supporting that these mutations lead to loss of function that impacts cellular fitness (Fig. S2, Table S4).

In summary, the screens had strong discriminatory power for gRNAs targeting regions of the genome that are required for normal cell proliferation and showed significant depletion or enrichment of a subset of the SLiM-targeting gRNAs.

### Defining a SLiM dependency map

Next, we defined a SLiM dependency map by analyzing the changes in gRNA abundances in a motif-centric manner. We reasoned that SLiMs targeted by multiple significant gRNAs are more likely to be required for cellular fitness. In accordance with this, we observed that 62.3% of the significantly changing gRNAs were found in only 11.5% of the SLiMs, suggesting that a subset of SLiMs are affecting proliferation. We further filtered the predicted motifs using AlphaFold structural information to more stringently enrich for motifs residing in intrinsically unstructured regions^19^. Next, we categorized the remaining 6,489 SLiMs into three classes of motifs: depleted motifs, which result in a proliferation defect upon mutation; enriched motifs, which provide a proliferation advantage upon mutation; and stable motifs, which show no observable effect upon mutation (examples in Fig. 2a-c). We applied three metrics to quantify essentiality and prioritize motifs by confidence across the four screens: the number of unique significant gRNAs per motif; the mean fold change of the gRNAs for each motif; and the distribution of these fold changes against the distribution of all gRNAs in the screen (Table S5). This data is made available through an interactive web resource for the community to explore SLiM function (https://slim.icr.ac.uk/projects/base_editing) (Fig. S3).

**Figure 2:**
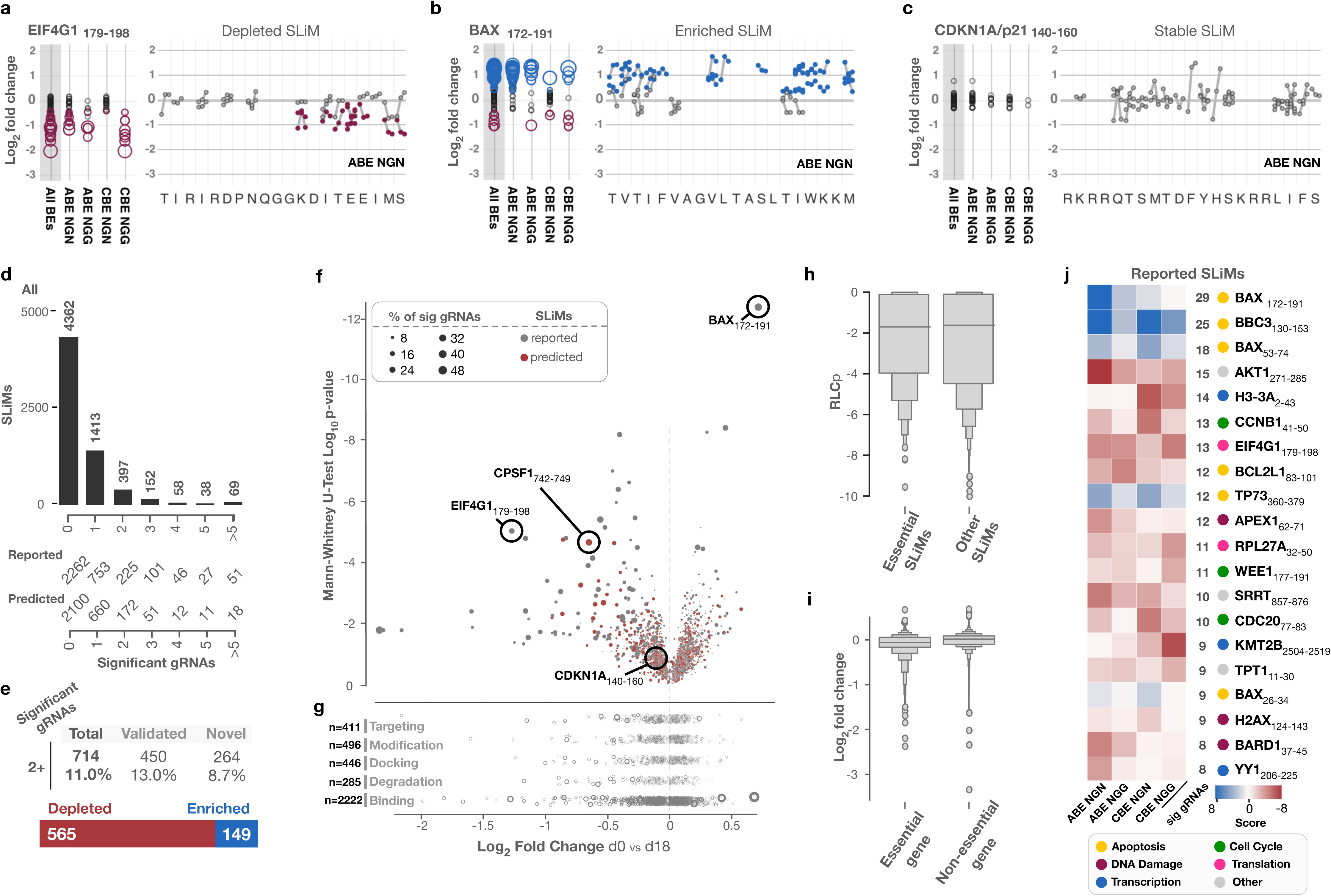
Identification of SLiMs required for optimal fitness. **a-c**, Examples of SLiM-containing regions with depleted (red) (**a**), enriched (blue) (**b**) or unchanged (grey) (**c**) gRNAs. Left panels: Y-axes indicate log_2_-fold changes of all gRNAs targeting the indicated motif summarised for the 4 screens (‘All BEs’) and individual screens. Circle size represents limma p-value. Right panels: Y-axes indicate the log_2_-fold change of a given gRNA predicted to target a given amino acid residue within the represented motif (indicated on the x-axis), with connected dots indicating the same gRNA edit across the biological triplicate of the ABE_NGN_ screen. In case gRNAs are predicted to target multiple residues, the log2 fold changes of the guide is shown for each residue. **d**, **e**, Number of SLiMs targeted by the indicated significantly enriched or depleted gRNAs. **f,** Mann-Whitney p-value versus log_2_ fold change using the most significant p-values for each motif from the four different screens. Red: *reported motif set*. Grey: *predicted motif set*. Size represents the percentage of significant gRNAs for each motif. **g**, Subdivision of motifs in panel **f** by ELM motif classes. **h**, Conservation of motifs categorised as essential versus non-essential. *RLCp*: SLiMPrint relative local conservation probability^35^. **i**, Log_2_-fold change of motifs categorised by HAP1 gene essentiality. **j**, Top enriched (blue) and depleted (red) reported SLiMs in the four screens. *Score:* log10 Mann-Whitney p-value, based on fold change, score is negative for depleted motifs, positive for enriched motifs. Circle color denotes general function of the motif-containing protein.

Overall, across the four screens, we identified 714 SLiMs in 626 proteins targeted by at least two unique gRNAs with significant depletion or enrichment (Fig. 2d). We consider these motifs the hits of our screen and will refer to them as ‘essential SLiMs’, that is SLiMs which affect normal cell proliferation when mutated. In contrast, the majority of motifs (4,362; 67.2%) were targeted solely by gRNAs that remained stable through the screening (Fig. 2d). We consider these to be non-essential for proliferation, although we cannot exclude the lack of phenotype to be due to low editing efficiency (see discussion). Of the essential SLiMs, 565 were depleted and 149 enriched upon mutation and 450 were from the *reported motif set* while 264 were from the *predicted motif set* (Fig. 2e-f).

We next categorized the *reported motif set* into high-level functionalities (Fig. 2g) and observed a small increase in essentiality for the binding (7.4%) and degradation (8.0%) classes of motifs compared to the modification, localization, and docking motif classes (5.8%, 6.0%, and 5.8%, respectively). We noted that many highly conserved SLiMs are not essential for cell proliferation and, contrary to expectation, that essential SLiMs are not more highly conserved than non-essential SLiMs (Mann-Whitney p-value 0.53, Fig. 2h, Table S5). Finally, as expected, when comparing SLiM essentiality to gene essentiality defined by HAP1 gene knockout screens^20^, essential motifs are more likely to be encoded in essential genes (Mann-Whitney p-value 1.69x10^-27^, Fig. 2i). However, we did observe essential motifs in proteins encoded by non-essential genes including the instance in Phosphatidate phosphatase LPIN1 (encoded by *LPIN1*) WLWGELP_273-279_ and Tyrosine-protein phosphatase non-receptor type 1 (PTP1B encoded by *PTPN1*) EVRSR_369-373_ investigated later.

Several of the SLiMs from the *reported motif set* with the highest number of significant gRNAs in the screen are in proteins with clear links to core biological functions (Fig. 2j). These motifs include the polyadenylate-binding protein-1 (PABP) binding motif in eIF4G required for the expression of polyadenylated mRNAs^21^ and the APC/C degradation motif (D box) of Cyclin B1 (CCNB1) required for mitotic exit^22^. Mutations in motifs with anti-apoptotic functions in many cases resulted in reduced cell proliferation, for example, mutations to motifs in Apoptosis regulator Bcl-2 family member BCL2L1^23^. Conversely, mutation of the BH3 motif in Apoptosis regulator BAX (BAX) and tetramerization motif in the Tumor protein p73 (TP73), both of which have pro-apoptotic functions^24–26^, resulted in a proliferative advantage. We performed a Reactome pathway analysis of the proteins harboring the essential SLiMs of the *reported motif set* compared to all proteins in the set. This showed an enrichment in essential motifs in pathways related to core biological processes such as transcription, RNA processing, cell cycle and apoptosis (Fig. S4a, Table S6). For the *predicted motif set*, functions were more diverse with no enrichment in any Reactome pathway. Mapping essential SLiMs to reported protein complexes retrieved from the Complex Portal^27^ suggested that numerous key complexes require multiple cooperative motif-mediated interactions for their function (Fig. S4b-d, Table S7). Three complexes stood out due to the number of essential motifs with five each: the Mitotic Checkpoint Complex (motifs in BUBR1 and CDC20) (Fig. S4b); Histone-lysine N-methyltransferase complex SET1A variant (motifs in SETD1A, RBBP5 and ASH2L) (Fig. S4c) and pre-mRNA cleavage and polyadenylation specificity factor complex (motifs in CPSF1, FIP1L1 and WDR33) (Fig. S4d). The latter two are of particular interest as they include several uncharacterized, yet essential motifs.

In summary, we used the base editing screens to define a dependency map of the SLiM-mediated interactome, identifying reported and predicted SLiMs affecting cell proliferation.

### gRNA behavior and editing outcomes are maintained across cell lines

To further confirm beyond the CPSF1 motif (Fig. S2) that introduction of the predicted edits by the base editor cause the observed effects on cellular fitness, we focused on the 714 essential motifs defined in the SLiM dependency map for validation. A major limitation of base editing screening approaches is that base editing outcomes are not measured in the screen but requires tedious post-screen validation experiments to confirm editing events (Fig. S2). During the preparation of this work, several base editing activity ‘sensors’ were published^28–31^ in which a surrogate target site is linked to the gRNA construct in the lentiviral vector, allowing for indirect measurement of base editing outcomes by NGS of the sensor (Fig. 3a). We thus designed a SLiM validation base editing library using a gRNA-sensor set-up (Fig. 3a-b). We focused on ABE_NGN_ due to its high performance with the relaxed NGN PAM requirement (Fig. 1h) and designed 8505 gRNA-sensor constructs targeting the 714 essential motifs defined in the SLiM dependency map, including both significant and non-significant gRNAs from the previous ABE_NGN_ screen (Table S8). In addition to positive and negative controls, eight gRNAs per SLiM-containing gene were designed to target splice sites to generate knockout-like phenotypes, allowing us to compare the effects of residue-level to gene level perturbations.

**Figure 3:**
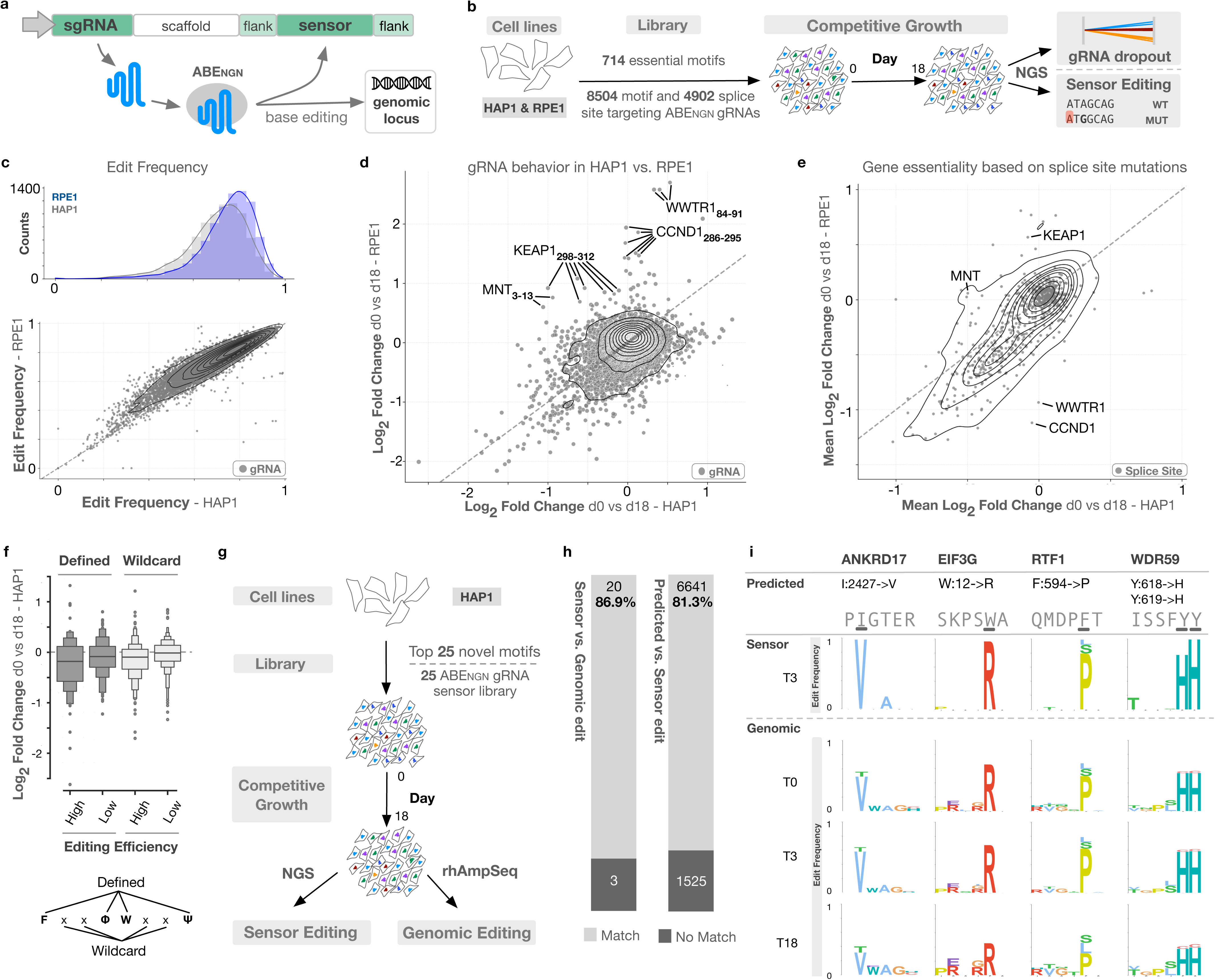
Validation of gRNA performance and editing outcomes. **a**, Schematic of the base editing sensor setup. **b**, Schematic of the experimental pipeline of the sensor screen targeting the 714 essential motifs and 8 splice sites per motif-encoding gene in HAP1 and RPE1 p53-/cells. **c**, Comparison of sensor editing frequencies in HAP1 and RPE1 cells. **d**, Comparison of SLiM-targeting gRNA log_2_ fold changes for the HAP1 and RPE1 cells. **e**, Comparison of mean log_2_ fold changes for gRNA sets targeting splices sites of SLiM-encoding genes for the HAP1 and RPE1 cells. **f,** Log₂ fold changes of gRNAs classified based on the ELM resource^5^ consensus as *defined* (specific, binding determining residues) or *wildcard* (flexible positions denoted as ‘x’) for significantly changing motifs (log₁₀ Mann–Whitney p < 0.01). gRNAs were further subdivided in the 50% highest and 50% lowest sensor editing efficiencies. **g**, Schematic of the experimental pipeline of the rhAmpSeq screen targeting the 25 predicted essential motifs with the top scoring gRNA for each, reading out sensor editing and genomic editing by NGS after pooled rhAmpSeq amplification of the 25 SLiM-containing loci. **h,** Comparison of the predicted or observed protein level edits between sources of edit information: *predicted*, based on any predicted edit for the ABE optimal editing window; *sensor*, based on the most common single edit in the sensor; and *genomic*, based on the most common single edit in the rhAmpSeq amplification of the genomic locus. **i,** Comparison of the *predicted* edit*, sensor* edit and *genomic* edits at day 0, 3, and 18 for four targets in the rhAmpSeq screen. Logos show the protein level edit frequency for the *sensor* and *genomic* data.

To evaluate the reproducibility of the HAP1 screen across cell types, the sensor screen was performed both in HAP1 cells and in a non-transformed RPE1 p53-/cell line, which we have previously used for base editing screening^32^. Cells were transduced with the lentiviral library at low multiplicity of infection and high gRNA coverage (>1000x) and allowed to grow competitively for 18 days. gRNA abundance and sensor editing in genomic DNA at day 0, 3 and 18 post selection were quantified by NGS (Fig. 3b), and changes in gRNA abundances from day 0 to day 18 were analyzed as for the primary screen (Table S9-S10). Editing outcome and frequency on the sensor in the starting population was evaluated at day 3 as this timepoint had higher editing frequencies than day 0 (Fig. 3c, Fig. S5a-b). Positive and negative controls behaved as expected, and no threshold was set on minimal editing frequency of a gRNA since it did not substantially improve the discriminatory power of the screen (Fig. S5c-d). First, we compared the changes in gRNA abundance to the primary screen data with the same editor (ABE_NGN_) to confirm reproducibility across screens (Fig. S5e). The gRNA behaviors were highly correlated although changes were less pronounced in the sensor screen.

A comparison of HAP1 and RPE1 cells revealed highly similar sensor editing frequencies (Pearson correlation r = 0.93, p < 0.001), enabling a direct comparison of gRNA behavior (Fig. 3c). Similarly, the gRNA abundance changes were correlated (positive controls: r = 0.73, p < 0.001, SLiM-targeting: r = 0.41, p < 0.001), despite a large proportion of the motif-targeting gRNAs showing no change, suggesting gRNA behavior is generally reproducible across cell lines (Fig. 3d, Table S9). However, we also observed some notable differences between HAP1 and RPE1 cells including the enrichment of gRNAs targeting specific motifs (Fig. 3d). In RPE1 cells, gRNAs targeting the AMBRA1 degron of Cyclin D1 and the 14-3-3 binding site in WW domain-containing transcription regulator protein 1 (WWTR1) are more highly enriched than in HAP1 cells. Conversely, gRNAs targeting the Sin3 motif of the Max-binding protein MNT (MNT) and the nuclear export signal of Kelch-like ECH-associated protein 1 (KEAP1) are depleted in HAP1 but enriched in RPE1 (Fig. 3d, Fig. S5f). These differences between cell lines likely reflect variation in the essentiality of the corresponding genes. As the library included gRNAs targeting splice sites of the genes harboring the SLiMs, we were able to compare the effect of SLiM mutation to gene essentiality from ‘knockouts’ created by base edited splice sites (mean log_2_ fold change of eight splice sites), which correlates well with DepMap^33^ knockout gene effects (Fig. S5g-h). Strikingly, for Cyclin D1 and WWTR1, mutation of the SLiM produces the opposite effect of the gene knockout, likely reflecting the growth suppressing role of these SLiMs, where motif loss would result in stabilisation of Cyclin D1^34–36^ and activation of WWTR1^37^. The motif-targeting gRNAs for MNT and KEAP1 follow the same pattern observed for their respective splice site mutations (Fig. 3d-e, Fig. S5f). Indeed, there was a trend for the log fold changes of significant motifs to align with the splice site-targeting gRNAs, confirming that motif essentiality tends to follow overall gene essentiality (Fig. S5i-j, Table S10).

Next, we looked more closely into reported SLiMs where the conserved key binding determinants (defined positions) are well-characterized to determine if mutations in these positions were more likely to cause a fitness defect than non-conserved flanking positions (wildcard positions) (Fig. 3f). We considered only the motifs which were significantly changing (Mann-Whitney p-value < 0.01, Table S10) and further separated the gRNAs into the 50% lowest and 50% highest editing. We observed that gRNAs targeting defined positions are more likely to cause a fitness effect than gRNAs targeting wildcard positions and that this effect is more prominent with high editing gRNAs than low editing gRNAs (Fig. 3f). This is consistent with the fitness effect being caused by disruption of SLiM-mediated binding through mutation of defined key binding residues.

Next, we wanted to further validate our top hits by probing editing in their genomic loci and track mutations during competitive growth. Here, we chose the top-scoring ABE_NGN_ gRNA for the top 25 predicted motifs and cloned a lentiviral gRNA-sensor library, which were transduced into HAP1 cells, followed by competitive growth and NGS of the sensor as described above (Fig. 3g). In addition, we performed pooled amplification and NGS of the 25 genomic target loci using rhAmpSeq^TM^ to obtain detailed information about editing at the genomic loci (Fig. 3g, Table S11, see also methods section). Inspecting the sensor and rhAmpSeq editing data, we observed highly similar outcomes across predicted, sensor-derived, and genomic locus edits (Fig. 3h, Fig. S6a-b). Exceptions included Eukaryotic peptide chain release factor GTP-binding subunit ERF3A (encoded by *GSPT1*) where K87E rather than the predicted K87G was the more predominant sensor and genomic locus edit and Kinase suppressor of Ras 1 (*KSR1*) where predicted and sensor editing matched (S453P) but the most predominant genomic locus edit was S453Y (Fig. S6a). In most cases, we observed that the relative frequency of the protein-level mutations changed from early to late timepoints (Fig. 3i, Fig. S6a). For example, ANKRD17 I2427V and EIF3G W12R were depleted over time, indicating that cells with these mutations were outcompeted from the population, consistent with the depletion of the respective gRNAs in the screens.

Together, these results demonstrate that i) the screen results were reproducible across two cell lines, ii) differences between the cell lines can largely be explained by differences in gene essentialities, iii) predicted, sensor, and genomic locus editing were highly similar, and iv) depleted gRNAs from the screen introduce mutations in the genomic target loci which are outcompeted over time.

### Base editing of a novel SLiM in CPSF1 links proliferation defects to loss of CSTF3 binding

Next, to validate that the introduction of the predicted edits by the base editor caused the observed effects on cellular fitness through disruption of SLiM-mediated interactions, we first focused on one of the top scoring SLiM from the *predicted motif set*, DDEEEMLYG_741-749_ in CPSF1 (Fig. 4a, Table S5). This SLiM shares significant similarity with the Cleavage stimulation factor subunit 3 (CSTF3)-binding motif in FIP1L1^38^, that is also a strongly depleted motif in our screen, particularly at core LYG and N-terminal acidic residues (Fig. 4a). We will refer to it as the LYG motif from here on. First, to explore whether mutations in this motif would lead to loss of binding partners, we transiently transfected HeLa cells with YFP-tagged full length CPSF1 wild-type or a mutant with alanine substitutions in the two amino acids that were mutated in the most depleted gRNAs targeting the motif, E743 and E744 (Table S3). The constructs were affinity purified (AP) from HeLa cells and a quantitative interactome comparison was performed using mass spectrometry (MS)-based proteomics (we will refer to this approach as AP-MS). This revealed that the CPSF1 LYG motif is required for binding to Cleavage Stimulation Factor complex subunits (CSTF1 and CSTF3), confirming that the LYG motif in CPSF1 presents another instance of a CSTF3 binding motif (Fig. 4b, Fig. S7a, Table S12). Consistent with this, an AlphaFold3 (AF3) prediction modelling of the full length version of the CPSF1-WDR33-CPSF4-RNA-CSTF3 complex positions the acidic LYG motif of CPSF1 in the basic SLiM-binding pocket on CSTF3 that was reported to bind the FIP1L1 motif^38^ (Fig. 4c, Fig. S7b).

**Figure 4:**
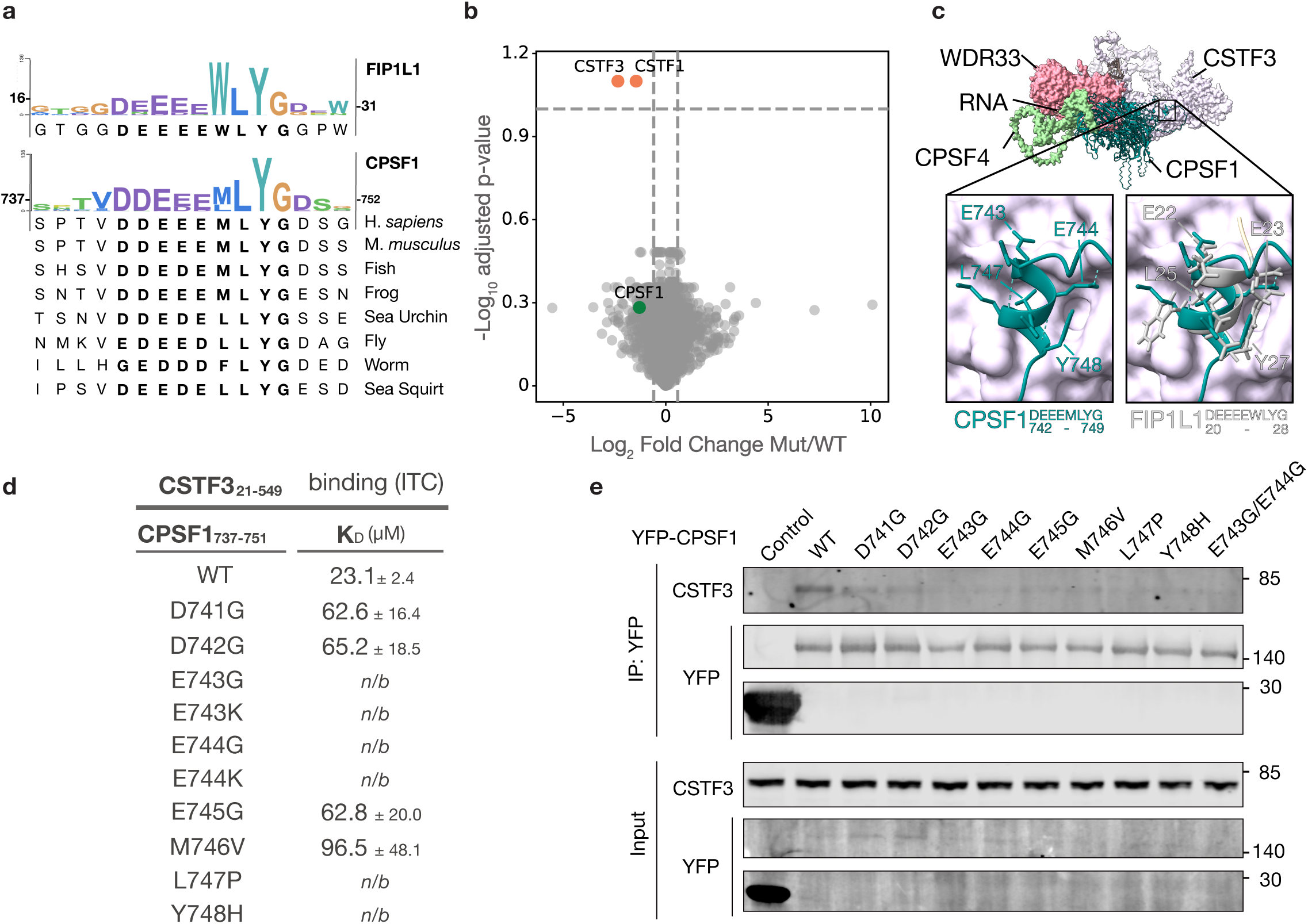
Validation of an essential predicted CSTF3 interaction motif in the cleavage and polyadenylation specificity factor CPSF1. **a**, Evolutionary conservation of the CPSF1 LYG (DDEEEMLYG_741-749_) motif and the CSTF3-binding FIP1L1 motif. **b**, Volcano plot showing mass spectrometry analysis of YFP-tagged full length CPSF1 WT vs E743A/E744A mutant (Mut) affinity purified from HeLa cells (n=4). The dashed lines indicate fold-change (1.5) and p-value (0.1) significance cut-offs. **c**, AlphaFold3 (AF3) prediction modelling the CPSF1-WDR33-CPSF30-RNA-CSTF3 complex (PTM score = 0.67; iPTM score = 0.74). Top panel: CPSF1 is presented as cartoon, the other proteins are represented as surface. Bottom left panel: detail of CPSF1_742-749_ (teal, cartoon representation) binding to CSTF3 (light purple, surface representation). Bottom right panel: detail of the structural alignement of CPSF1-WDR33-CPSF30-RNA-CSTF3 AF3 model with the FIP1L1-CSTF3 structure (PDB: 7ZY4), showing FIP1L1_20-28_ (grey, cartoon representation) binding to the CPSF1-binding pocket of CSTF3 (light purple, surface representation). **d**, ITC binding data of CPSF1_737-751_ wild-type (WT) and mutant peptides to CSTF3_21-549_. **e**, Western blot analysis of full length WT and mutant YFP-CPSF1 affinity purifications from HeLa cells probed for CSTF3 and YFP, representative of two independent experiments.

To directly link the observed effects on cellular fitness of the base editor-induced mutations in CPSF1 to the loss of binding to CSTF3, we tested the impact of 10 predicted single amino acid mutations achievable by base editing on CSTF3 binding. Isothermal titration calorimetry (ITC) affinity measurements of CPSF1 peptides to recombinant CSTF3_21-549_ revealed reduced binding of all mutants compared to wild-type (Fig. 4d, Fig. S8, Table S13). These data were consistent with the co-affinity purification of CSTF3 with wild-type but not mutant full length YFP-tagged CPSF1 from HeLa cells (Fig. 4e). The affinity measurements revealed that the ABE mutations introduced at E743, E744, L747 and Y748 had the largest impact on binding, consistent with the key buried residues in the reported structure of the FIP1L1 LYG motif bound to CSTF3^38^ (Fig. 4c, Fig. S7b).

Collectively, these data demonstrate that single amino acid mutations introduced into a SLiM by base editing are sufficient to disrupt protein interactions resulting in a fitness defect that can be measured by proteome-wide screening.

### Essential predicted motif instances largely represent novel motif families

We next investigated the remaining 263 essential SLiMs from the *predicted motif set* to determine if they belong to known or novel motif families (Fig. 5a for top scoring motifs and Table S5). We first compared these SLiMs to the specificity determinants of previously characterized domain-binding motifs (see method section), revealing 39 predicted instances with significant similarity to existing SLiM families (Fig. 5b, Table S14). The majority of these examples belong to well characterized motif classes. For example, the ERTPLL_306-311_ motif in the E3 ubiquitin-protein ligase RNF167 (RNF167) strongly resembles an Adaptin binding Endosome-Lysosome-Basolateral sorting signal (Fig. 5c), aligning with the role of RNF167 in regulating endosome trafficking^39^. Another example is the SWADQVE_11-17_ motif in Eukaryotic translation initiation factor 3 subunit G (EIF3G), which had significant similarity to the binding determinants of a motif binding to MIF4GD. The motif class had previously only been studied in fish, with MIF4G domain-binding motifs in *Danio rerio* Stem-loop binding protein (SLBP) and mRNA-export factor DBP5 (DBP5)^40^. Therefore, the predicted EIF3G instance would extend the taxonomic range of the motif to human proteins (Fig. 5d). To test this, we again used an AP-MS approach comparing full length YFP-tagged human EIF3G wild-type to a mutant with the amino acids of the predicted motif substituted with alanine residues. This showed strong and specific binding to human MIF4G that was lost upon mutation of the SLiM, consistent with the observations in *Danio rerio* (Fig. 5e). Next, we compared the binding determinants of the 264 predicted essential motifs to each other and observed only 16 instances with similarity to other predicted motifs. Those that shared binding determinants consisted of motifs in paralogous proteins, proline-rich motifs, and a cluster of transcription factors that shared hydrophobic motifs resembling transactivation domain^41^ (Table S14). Strikingly, the specificity determinant analysis indicated that 225 predicted essential SLiMs showed no significant similarity to any known motif class, suggesting that the majority of the predicted SLiMs do not belong to existing motif families (Fig. 5b, Table S14).

**Figure 5:**
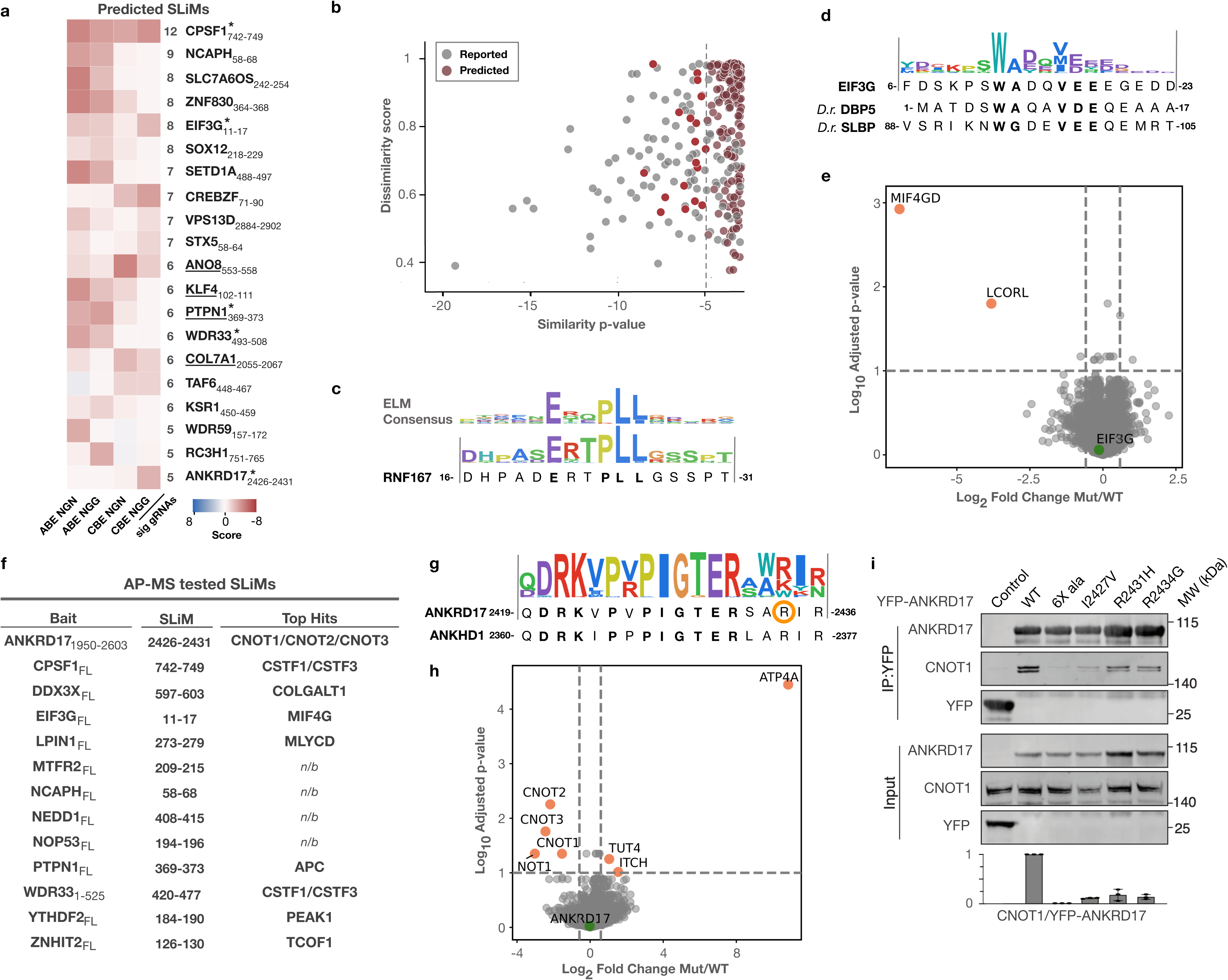
Characterisation of essential predicted motifs. **a**, Top essential predicted SLiMs in the four screens. *Score:* log10 Mann-Whitney p-value, based on fold change. Total number of significant gRNAs are indicated. **b**, Comparison of the essential motifs to the specificity determinants of the ELM resource^5^ using CompariPSSM^57^. *Similarity p-value*, quantifies the similarity of the binding determinants. *Dissimilarity score*, quantifies key binding determinants with strong dissimilarity. Bright red dots correspond to motif pairs with with similarity scores above the default CompariPSSM cutoff. **c**, Conservation logo of the ELM Adaptin-binding Endosome-Lysosome-Basolateral sorting signals class (TRG_DiLeu_LyEn_5) and the conservation logo of RNF167 motif ERTPLL_306-311_ **d**, Conservation logo of EIF3G motif SWADQVE_11-17_ and *Danio rerio* DBP5 and SLBP5 MIF4G binding motifs. **e**, Volcano plot showing affinity purification coupled to mass spectrometry (AP-MS) analysis of YFP-tagged full length EIF3G WT vs motif mutant (Mut; AAAAAAA_11-17_) purified from HeLa cells (n=4). The dashed lines indicate fold-change (1.5) and p-value (0.1) significance cut-offs. **f,** Summary of AP-MS analysis of YFP-tagged full length bait proteins (except for ANKRD17_1950-2603_ and WDR33_1-525_ where fragments were used) purified from HeLa cells. Top significantly lost binder or, if relevant, top lost complex is shown, comparing WT to motif mutants. n/b, no binder. **g**, Conservation logo of ANKRD17 PIGTER_2426-2431_ motif with surrounding sequence and homologous region of ANKHD1. R2434 mutated in Chopra-Amiel-Gordon syndrome is circled. **h**, Volcano plot of AP-MS for motif-mutant (Mut; AAAAAAA_2425-2431_) versus WT YFP-ANKRD17_1950-2603_, purified from HeLa cells (n=4). The dashed lines indicate fold-change (1.5) and p-value (0.1) significance cut-offs. **i**, Representative Western blot and quantification of affinity purified YFP-ANKRD17_1950-2603_ WT and mutant from HeLa cells, probed for CNOT1 and YFP (n=3, error bars represent mean +/standard deviation).

To further explore the predicted essential motifs, we expanded our AP-MS approach, uncovering putative binding partners for seven of the additional 11 baits tested (Fig. 5f, Fig. S9, Table S12). In WDR33, the proliferation screen had uncovered an essential IPG motif (IPGMG_441-445_), and further inspection of this region revealed an extra conserved copy of this consensus motif (IPGLDWG_471-477_) as well as an additional LPGM_420-423_ motif (Fig. S10a). Mutation of all three resulted in the loss of binding to the Cleavage Stimulation Factor subunits CSTF1 and CSTF3 in AP-MS (Fig. S9). Direct binding of the IPG motifs in WDR33 to CSTF3 was validated by pull-down of CSTF3_21-549_ with a WDR33 fragment covering both IPG motifs, WDR33_431-485_, and mutation of both motifs disrupted binding (Fig. S10b). This unstructured region of WDR33 would be in close proximity to CSTF3 based on the structure of the CPSF1-WDR33-CPSF4-RNA-CSTF3 complex (6URO) which contains the domain directly N-terminal of the IPG motifs^42^, and our results suggest that the IPG motifs help stabilise the interaction. This possibility is supported by AF3 predictions using full length proteins, where the essential WDR33_441-444_ motif was confidently modelled in a binding pocket in CSTF3 (Fig. S10c). Another interesting observation was the loss of binding between the tyrosine phosphatase PTP1B and Adenomatous Polyposis Coli (APC) protein upon mutation of the EVRSR_369-373_ motif in PTP1B (Fig. S9, Table S12). This novel interaction suggests that the actions of PTP1B and APC in regulating β−catenin during Wnt signaling might be coordinated^43^. This is an example of an essential SLiM in a non-essential protein, where mutation leads to a fitness defect, possibly through the creation of a dominant negative version of the protein causing misregulation of β−catenin.

### A disease linked mutation in ANKRD17 disrupts SLiM binding to the CCR4-NOT complex

We next focused on whether we could obtain novel mechanistic insight into disease-causing mutations potentially affecting SLiM function. Of the 714 essential SLiMs, 104 had reported pathogenic disease mutations (extracted from PepTools^44^) within the motif or in the immediate flanking regions (3 amino acids on either side) with 57 of these being point mutations similar to the outcome of base editing (Table S15). We focused our attention on a PIGTER_2426-2431_ motif present in the mRNA binding^45^ protein Ankyrin repeat and KH domain-containing 17 (ANKRD17), representing a novel uncharacterised motif family (Fig. 5g). The rhAmpSeq screen confirmed that mutation of the SLiM (I2427V) in the genomic locus resulted in depletion from the cell population over time, consistent with a fitness defect (Fig. 3i). An additional instance of the motif in the ANKRD17 paralogue Ankyrin repeat and KH domain-containing 1 (ANKHD1), mapped in the specificity determinant analysis (Table S14), was also found to be essential in our screen. In ANKRD17, a disease mutation (R2434->G) of unknown consequence was present C-terminal to the motif, PIGTERSA**R**_2426-2434_. This mutation is classified as likely pathogenic in ClinVar, linked to Chopra-Amiel-Gordon syndrome (CAGS), a neurodevelopmental syndrome caused by heterozygous loss of ANKRD17 function^46^. Using our AP-MS approach, we found that upon mutation of the entire motif in ANKRD17 binding to CNOT1, CNOT2, and CNOT3 was lost, suggesting that the PIGTER motif mediates binding to the NOT module of the CCR4-NOT deadenylase complex involved in mRNA regulation, consistent with published proximity mapping data^47,48^ (Fig. 5h). We validated that the predicted amino acid mutations introduced by the base editors in ANKRD17 disrupted the binding to CNOT1 by affinity purification from HeLa cells, and importantly, we confirmed that the R2434G CAGS disease mutation disrupted binding to CNOT1 (Fig. 5i). We speculate that the defect in ANKRD17 function in patients with this mutation is due to deregulation of specific mRNAs bound by ANKRD17 that are no longer channeled to the CCR4-NOT complex^46^.

## Discussion

SLiMs constitute one of the most abundant regulatory elements in unstructured regions of the human proteome and can readily arise *ex nihilo* or be disrupted through disease mutations. Until now, our understanding of SLiM contribution to cell function has been limited due to lack of efficient endogenous functional screening methods. In this study, we utilize base editor screening to create a comprehensive map of cellular SLiM dependency in HAP1 cells and show that mutations introduced by base editors are sufficient to disrupt SLiM mediated interactions, allowing us to couple disruption of interactions to cellular fitness effects. Importantly, the interpretation of our data is not confounded by structural considerations as we target SLiMs in unstructured regions. Our screens reveal that numerous motifs are required for optimal cell proliferation both in HAP1 and RPE1 cells. By defining a set of essential SLiMs, we provide a priority list for future studies, as well as gRNAs to specifically target these. We provide a user-friendly web-based resource (https://slim.icr.ac.uk/projects/base_editing) that allows for interactive exploration of our comprehensive dataset, thereby providing a valuable tool for the research community. We furthermore show that binding partners of essential SLiMs can be identified by a streamlined AP-MS approach comparing affinity-tagged wild-type protein to a variant where the SLiM is mutated. This approach can now be applied to many of the predicted SLiMs we identify as essential, followed up by mutational binding analyses and modelling to characterize mode of binding.

While we cannot exclude that some motifs do not score due to low editing efficiency of the gRNAs targeting them, our sensor screen suggests that the majority of gRNAs, at least for ABE_NGN,_ are editing efficiently (Fig. 3c). We therefore favor that the ∼90% non-scoring motifs are not required for proliferation in HAP1 cells under normal cell culture conditions. Importantly, our data show that essential SLiMs are more likely to occur within essential proteins (Fig. 2j), arguing that such fitness screens are mainly able to measure mutational effects in essential proteins. Thus, we consider the non-scoring motifs a large unexplored SLiM space with context specific functions. Our gRNA libraries now provide a powerful tool to explore these context specific functions of SLiMs by applying the libraries on additional cell types or during cell perturbations. Our comparison of RPE1 and HAP1 cells already confirms the utility of our libraries to identify cell line specific differences in SLiM dependencies. A further avenue to explore is to employ the gRNA libraries under conditions where cells are treated with therapeutic agents, potentially identifying SLiM interactions that can constitute novel vulnerabilities that can be exploited for drug development^49^.

In conclusion, our work highlights that base editing technologies can be applied to discover novel protein binding regions and prioritize interaction interfaces based on their importance to cell fitness. Our approach provides a blueprint for future efforts to further dissect SLiM function.

## Acknowledgments

Work at the Center for Epigenetic Cell Memory, Danish Cancer Institute is supported by The Danish National Research Foundation (DNRF195). Work at the Novo Nordisk Foundation Center for Protein Research is supported by NNF14CC0001. Work in the Nilsson lab is supported by grants from the Novo Nordisk Foundation (NNF0082227 and NNF0065098), DFF-FNU (4283-00188B) and the Danish Cancer Society (R269-A15586-B71). N.E.D. is funded by a Cancer Research UK Senior Cancer Research Fellowship (C68484/A28159). E.P.T.H. is funded by a Lundbeck Foundation Postdoc Grant (R380-2021-1221). We thank the protein production and characterization facility at Novo Nordisk Foundation Center for Protein Research for all their help with the project. We thank the Genomics Platform at Novo Nordisk Foundation Center for Protein Research and Center for Stem Cell Medicine (reNEW) for technical support and use of instruments. NGS data processing and analysis were performed using the DeiC National Life Science Supercomputer at Technical University of Denmark (www.computerome.dk).

## Methods

### SLiM library design

SLiM-containing regions were collected from three sources (time of download: April 2022): 1) manually *curated* motif information and 2) motif-containing *structure* processing to retrieve experimentally validated motifs (as a source for the *reported* dataset); and 3) *evolutionary footprinting* to define putative motifs (source for the *predicted* dataset). *Curated* data was extracted from the motif literature, or retrieved programmatically from the ELM datasets. *Structural* data on motif instances was programmatically extracted from PDB by screening all PDB structures that contain a biological assembly with 2+ chains; defining disordered peptides in these structures using FLIPPER^50^; and extracting SLiMs using the discriminator of bound footprint <1500 Å^2^, mono-partite interface as defined by a single distinct secondary structural properties in the bound conformation, and defined by FLIPPER as disordered across the entire chain. *Evolutionary footprinting* data was defined using the SLiMPrints algorithm to define conserved regions with SLiM-like evolutionary patterns in the IDRs of the human proteome^51^. Significant predicted motifs were defined with two cut-offs to enrich for functionally important motifs, for instances in DepMap essential proteins with mean gene effect < -0.5 or overlapping disease relevant SNVs, a SLiMPrints significance cutoff of 0.005 was used, otherwise a cut-off of 0.0005 was applied. Accessibility filter was then applied to the SLiMPrints prediction to enrich for accessible intracellular motif instances by filtering predictions with an IUPred score less than 0.33 and removing peptides overlapping Transmembrane, Extracellular or Lumenal annotation in UniProt^52,53^.

### gRNA library design

gRNAs were designed to target the protein level of the motifs in the SLiM library based on the PAM (NGG or NGN), editing window (nucleotides 4-8 in the target DNA of the gRNA for all editors) and editing preference (ABE A→G, CBE C→T) of each base editor applied in the screen (ABE_NGN_, CBE_NGN,_ ABE_NGG_, and CBE_NGG_). The UniProt defined protein region was mapped to the corresponding transcript in Ensembl. The genomic sequence information of the gene, the oligonucleotide sequence of the gene in the genome, the CDS (CoDing Sequence) of the transcript, and the intronless oligonucleotide sequence product of the gene that is translated into a protein, were retrieved from Ensembl. The position of the target nucleotides in the CDS were determined by mapping the UniProt start and stop positions to the CDS, and the fidelity of the mapping was confirmed by mapping the peptide sequence to the translated CDS of the region. In the cases where no translated Ensembl transcript maps to UniProt protein sequence, the longest Ensembl transcript was retrieved, the target peptide region was mapped to the longest translated Ensembl transcript and the target positions are adjusted to account for the remapping. Next, the Ensembl CDS and genomic sequence were aligned and the target genomic sequence nucleotides for editing were determined through the translated Ensembl CDS. This step allows for editing of nucleotides flanking intron-exon boundaries. Potential PAM sites upstream and downstream of the target genomic sequence nucleotides were identified on both strands. PAM sites that positioned the edit window of the base editor over the target genomic sequence nucleotides were investigated further. The editing window was searched for nucleotides corresponding to the preference of the base editor to find potential nucleotide edits. If one or more editable nucleotides were available for a window all edits were made simultaneously *in silico* and the resulting protein-coding changes were compared to the wild-type protein sequence. gRNAs predicted to result in non-synonymous mutations were retained as part of the SLiM dependency map gRNA library. A set of previously described control gRNAs^16^ including 62 positive control targeting splice sites of essential genes and 255 negative controls targeting intergenic regions of the genome and non-targeting gRNAs were also included in the library design.

### Library cloning

Each gRNA in the library was extended to create a pool of oligonucleotide sequences (83 nt) following a previously published design^16^. The oligonucleotide consisted of primer sites for differential amplification, flanking appropriate overhang sequences with Esp3I restriction enzyme recognition sites and the gRNA: 5’-[Forward primer, 20 nt]CGTCTCACACCG[sgRNA, 20 nt]GTTTCGAGACG[Reverse Primer, 20 nt]. The oligonucleotide pool was synthesised by GenScript (CustomArray). Individual subpools were amplified in a reaction of 25 µl 2x NEBnext PCR master mix (New England Biolabs), 1 µl oligonucleotide pool (10 ng/µl), 2.5 µl each of forward and reverse primer at a final concentration of 0.5 uM, and 19 µl water, using a unique primer combination for each subpool (Table S16).

The number of PCR reactions per subpool was determined by scaling to 24 reactions per 92,000 gRNAs. PCR cycling conditions were: (1) 98 °C for 30 s; (2) 53 °C for 30 s; (3) 72 °C for 30 s; (4) go to (1) x 19; (5) 72 °C for 60 s; (6) 10 °C hold. Amplicons were PCR-purified (NucleoSpin Gel and PCR Clean-up, Macherey-Nagel). Library vector was pre-digested with FastDigest Esp3I (Thermo Fisher Scientific) and gel-purified (NucleoSpin Gel and PCR Clean-up, Macherey-Nagel). The following library vectors were used: CBE_NGG_: pRDA_256 (BE3.9Max-SpCas9); CBE_NGN_: pRDA_478 (BE3.9Max-SpG); ABE_NGG_: pRDA_426 (ABE8e-SpCas9); and ABE_NGN_: pRDA_479 (ABE8e-SpG). pRDA_256 (Addgene #158581), pRDA_478 (Addgene #179096), pRDA_426 (Addgene #179097), pRDA_479 (Addgene #179099) were gifts from John Doench & David Root. Amplicons were cloned into library vectors by Golden Gate reactions containing 25 µl 2x Rapid Ligase Buffer (Qiagen Enzymatics), 0.25 µl Bovine Serum Albumin (20 mg/mL) (New England Biolabs), 1 µl FastDigest Esp3I (1 U/µl), 0.25 µl T7 Ligase (3x10^6 U/mL) (Qiagen Enzymatics), 15 ng purified amplicon, 100 ng digested vector, and water in a total volume of 50 uL. The number of Golden Gate reactions performed per subpool was determined by scaling to 56 reactions per 92,000 gRNAs. Golden Gate reaction conditions were: (1) 37 °C for 5 min; (2) 20 °C for 5 min; (3) go to (1) x 24; (4) 10 °C hold. Ligated plasmid product was purified first by PCR-purification and then by isopropanol precipitation in the following reaction: 300 µl PCR-purified plasmid, 300 µl isopropanol, 3 µl GlycoBlue Coprecipitant (final conc. 75 ng/µl) (Thermo Fisher Scientific), 50 mM NaCl. Samples were vortexed (10 s full speed) and incubated 15 min at RT before centrifugation at 16,000xg for 15 min at RT. Precipitated DNA was washed twice with ice-cold 80% Ethanol, air-dried and resuspended in TE buffer (1 µl TE buffer per Golden Gate reaction) at 55 °C for 10 min. The number of electroporations per subpool was determined by scaling to 16 electroporations per 92,000 gRNAs. The purified plasmid library was electroporated into Endura Electrocompetent cells (Lucigen) and grown at 30 °C for 16 h on agar with 100 µl/mL carbenicillin. Colonies were scraped off and plasmid DNA was prepared (NucleoBond Xtra Maxi EF, Macherey-Nagel). To confirm library representation and distribution, inserts were amplified from plasmid library in a reaction of 25 µl 2x NEBnext PCR master mix, 1 µl plasmid library (10 ng/µl), 2.5 µl each of i5 and i7 primers at a final concentration of 0.5 uM, and 19 µl water, using unique primer combinations (Table S15). PCR cycling conditions were: (1) 98 °C for 30 s; (2) 98 °C for 10 s; (3) 55 °C for 30 s; (4) 65 °C for 15 s; (5) go to (2) x 19; (6) 65 °C for 5 min; (7) 10 °C hold. Amplicons were first gel-purified and then re-purified by PCR-purification. Samples were sequenced on a NextSeq500 or a NextSeq2000 (Illumina) with a 2% spike-in of PhiX with 21 (NextSeq500) or 22 (NextSeq2000) dark cycles. Fastq files were generated using bcl2fastq v.2.19.1 and reads were trimmed to 20 bp using cutadapt 1.18, removing a variable number of basepairs at the start and end depending on the size of the primer stagger. MAGeCK^54^ 0.5.8 was used to assign the trimmed reads to the gRNAs in the library and create the gRNA count matrix.

### Virus production

HEK 293T/17 cells (ATCC, CRL-11268) were seeded in DMEM GlutaMax (Gibco) + 10% FBS (Gibco) + 100 U/mL penicillin–streptomycin (Gibco). Medium was exchanged with Opti-MEM (Gibco) + 5% FBS 24h later for co-transfection of the base editing sgRNA library with lentiviral packaging plasmids pMD2.G and psPAX2 using Lipofectamine 3000 (Invitrogen). 6 hours after transfection, the medium was exchanged for DMEM GlutaMax + 10% FBS + 100 U/mL penicillin–streptomycin + 1% bovine serum albumin. 48 hours after transfection, viral particles were collected, filtered through a 0.45 μm syringe filter, and stored at -80°C. pMD2.G and psPAX2 were gifts from Didier Trono (Addgene plasmid # 12259; http://n2t.net/addgene:12259; RRID:Addgene_12259 and Addgene plasmid # 12260; http://n2t.net/addgene:12260; RRID:Addgene_12260).

### Competitive Growth Screen

HAP1 cells (#C859, Horizon Discovery, SNB: 35369) were cultured in IMDM GlutaMax (Gibco) supplemented with 10% FBS and 100 U/mL penicillin–streptomycin and passaged every three days by trypsinization in 0.05% trypsin-EDTA (Gibco). Each screen was performed as a triplicate with three separate transductions at a coverage above 675-fold sgRNA representation, which was maintained throughout the screen. HAP1 cells were transduced with the indicated lentiviral libraries at a low multiplicity of infection (MOI; 0.3-0.4) by treating cells with 8 μg/mL polybrene and lentiviral supernatants for 24 hours. Viral titers required to achieve an MOI of ∼0.3 were pre-determined in a titration experiment. Transduced cells were selected by treatment with 1.5 μg/mL puromycin for 48 hours. After selection, cells were cultured for 21 days. Genomic DNA was extracted from cell pellets harvested after completion of selection, which we consider the start of the screen (d0), and after 18 (d18), and 21(d21) days of culturing. ∼600 bp plasmid library fragments containing the sgRNA sequences were amplified from the genomic DNA by PCR using NEBNext Ultra II Q5 Master Mix with the LCV2_forward and LCV2_reverse primers (Table S16). A second PCR reaction introduced i5 and i7 multiplexing barcodes (Table S16), whereafter ∼220 bp PCR products were gel-purifed and sequenced on Illumina NextSeq2000 with a 2% spike-in of PhiX with 22 dark cycles. Fastq files were generated using bcl2fastq v.2.19.1 and reads were trimmed to 20 bp using cutadapt 1.18, removing a variable number of basepairs at the start and end depending on the size of the primer stagger. MAGeCK 0.5.8 ^54^was used to assign the trimmed reads to the gRNAs in the library (motif-targeting and control gRNAs (essential splice sites, non-targeting, intergenic)) and create the count matrix.

### Base editing data processing

NGS count data from the four screens (three timepoints and three replicates each) were processed to define significantly changing gRNAs using a limma statistical analysis^18^. Low-abundance gRNAs were removed (counts <30). The vast majority of gRNAs had a single exact match in the genome, however, non-specific gRNAs with off-target genome binding sites showed increased dropout correlated with the number of off-target binding sites. Consequently, gRNAs with 5 or more off-target matches in the genome were removed from downstream analysis, leading to the exclusion of 483 gRNAs. Raw sequencing counts were normalized for input into the limma software by dividing the number of reads for a specific gRNA by the total number of reads from the sample, multiplying by a million to the define the transcripts per million (TPM) and calculating the log_2_ of the gRNA TPM to define the log_2_ transcripts per million (log2TPM). The log2TPM values were compared using limma, resulting in fold change (FC) and p-values for each gRNA. These p-values were compared for the positive and negative controls, and a p-value cut-off of 0.01 showed a false positive rate of ∼1% across all four screens, with a true positive rate of up to 79%. Consequently, a p-value of less than 0.01 was set as the threshold considered to impact cell proliferation.

Next, the gRNAs mapping to each motif-containing region were analyzed to define groups of significantly changing gRNAs. The main metrics used was the number of unique significant gRNAs per motif across all screens. Per screen, the direction of the effect on proliferation was quantified as the mean fold change of the gRNAs to define depletion or enrichment, and the fold change of the gRNAs for a region were compared against the fold change for all gRNAs in the analysis using a Mann–Whitney U test to calculate an enrichment or depletion p-value. AlphaFold2 was applied as post-screen filtering to flag motifs that are buried or in regions incompatible with binding. The structural context of each region was defined using AlphaFold2 models. Full-length protein AlphaFold2 models were retrieved from the AlphaFold2 protein structure database (https://alphafold.ebi.ac.uk/). The structured regions of the proteins were extracted using the xProtCAS tool^55^ using a graph-based community detection algorithm on the AlphaFold2 predicted aligned error (PAE) matrix. Based on the xProtCAS, the determined structural state of each residue was classified as Domain, Loop or Unstructured. The unstructured proportion contains IDRs and coiled coils; coiled coil regions were defined as predicted helices greater than 25 amino acids in length; the remaining residues were classified as IDR. Potential intramolecular IDR Domain contacts were also determined as the first or last region in an autonomous structural module of length less than 30 connected by a loop of greater than 20 amino acids. The SLiMPrints portion of the initial library contained 760 predicted motif instances and 10,606 gRNAs that were filtered after AlphaFold2 predicted they were located in coiled coil or domain regions. As a result, 6,489 of the 7,293 motif-containing regions were retained for downstream processing and analysis.

### Region annotation

Regions were annotated using the PepTools peptide annotation tool (https://slim.icr.ac.uk/tools/peptools/) to define overlapping mutagenesis (source: UniProt), disease relevant SNVs (source: gnomAD, dbSNP, COSMIC curated, TOPMed, NCI-TCGA COSMIC, NCI-TCGA, ExAC, Ensembl, ESP, ClinVar, 1000 Genomes retrieved from the EBI API), and other features of interest (source: UniProt)^44^. Protein complex members were retrieved from the Complex Portal^27^. Pathway enrichment analysis was performed using the PANTHER enrichment tool^56^ to identify enriched Reactome pathway annotation using the PANTHER API (https://www.pantherdb.org/services/) by comparing essential motif-containing proteins to the protein set tested in the base editing analysis. The specificity determinant comparison analysis was performed with the CompariPSSM tool^57^. The specificity determinants of the regions targeted in the base editing analysis were extracted as conservation-derived PSSMs. An orthologue alignment was created for each region by running the GOPHER^58^ algorithm on the region-containing protein against a search database of metazoan proteomes. For each region, the columns corresponding to the edited region were extracted from the alignment and converted into a frequency PSSM. Frequency PSSMs were also created from aligned experimentally tested motif instances for motif classes from the ELM resource (http://elm.eu.org/) and a more extensive in-house curated MoMaP motif dataset (https://slim.icr.ac.uk/momap/). The base-edited PSSM set was then compared against the ELM PSSM dataset, MoMaP dataset, and itself using the CompariPSSM tool. Significant matches were defined as comparisons with a CompariPSSM similarity significance of <1x10^-5^, dissimilarity score of <1 and PSSM overlap >75%.

### Library design for the sensor and rhAmpSeq screens

The *sensor screen* library includes all gRNAs targeting the 714 motifs with 2+ unique significant gRNAs listed in Table S5, filtered to retain only gRNAs predicted to result in non-synonymous mutations with ABE_NGN_ (ABE8e-SpG; total 8505 gRNAs, Table S8). To generate the sensor region, each gRNA was mapped back to the genome, and a 41nt sequence overlapping the targeted site was extracted, consisting of an 11nt upstream flank, the 20nt gRNA target, and a 6nt downstream flank. Eight gRNAs predicted to mutate splice sites of each targeted gene were also included in the library. In addition to the positive and negative controls gRNAs from the original screen, the library included 500 positive control gRNAs targeting splice sites of essential genes and 500 negative controls targeting intergenic regions of the genome (250 gRNAs) and non-targeting gRNAs (250 gRNAs). The library for the *rhAmpSeq screen* constitues the top scoring ABE_NGN_ gRNA for the top 25 novel motifs in a gRNA-sensor set-up (Table S8). Each gRNA and sensor construct in the libraries were designed inspired by the general architecture from published work^28^ with an additional inclusion of a 4 nt barcode at the 5’ end of each insert^31^ to create a pool of oligonucleotide sequences. Each oligonucleotide consists of primer sites for differential amplification flanking appropriate overhang sequences with Esp3I restriction enzyme recognition sites and the gRNA, gRNA scaffold sequence, termination site, sensor site (gRNA with flanking sequences) and bar code (206 nt): [Fw_primer,20nt]CGTCTCACACCG[gRNA,20nt]GTTTGAGAGCTAGAAATAGCAAGTTCAAATAAGGCTAGTCCGTTATCAACTTGAAAAAGTGGCACCGAGTCGGTGCTTT TTT[flank,11nt][gRNA,20nt][flank,6nt][barcode,4nt]GTTTCGAGACG[Rv_primer,20nt].

### Sensor and rhAmpSeq screen library cloning

An oligopool comprising both *sensor* and *rhAmpSeq* screen libraries was ordered as a Sureprint OLS Hifi Oligo library 171-210 nt, 15K (Agilent). The libraries were amplified in a total of 30 reactions for the *sensor* library and 4 reactions for the *rhAmpSeq* library, each reaction consisting of 25 µl 2x NEBnext PCR master mix, 1 µl oligonucleotide pool (1 ng/µl), 0.25 µl each of forward and reverse primer at a final concentration of 0.5 uM, and 23.5 µl water, using a unique primer combination for each library (Table S16). PCR cycling conditions were: (1) 98 °C for 30 s; (2) 53 °C for 30 s; (3) 72 °C for 30 s; (4) go to (1) x 14; (5) 72 °C for 60 s; (6) 10 °C hold. Amplicons were PCR-purified. Library vector pLenti-U6-sensor-ABE8e-nSpG(D10A) was cloned by KpnI+EcoRI restriction of pRDA_479 and insertion of synthetic DNA carrying U6 promoter followed by BsmBI cassette, thus removing the gRNA scaffold sequence to allow insert of the gRNA-scaffold-sensor cassette. The vector was pre-digested with FastDigest Esp3I and gel-purified. Amplicons were cloned into pLenti-U6-sensor-ABE8e-nSpG by Golden Gate cloning in a total of 15 reactions for the *sensor library* and 6 reactions for the *rhAmpSeq library*. Each reaction contained 25 µl 2x Rapid Ligase Buffer, 0.25 µl Bovine Serum Albumin (20 mg/mL), 1 µl FastDigest Esp3I (1 U/µl), 0.25 µl T7 Ligase (3x10^6 U/mL), 15 ng purified amplicon, 100 ng digested vector, and water in a total volume of 50 µL. Golden Gate reaction conditions were: (1) 37 °C for 5 min; (2) 20 °C for 5 min; (3) go to (1) x 24; (4) 10 °C hold. Ligated plasmid product was purified first by PCR-purification and then by isopropanol precipitation as described above. The purified plasmid library was electroporated into Endura Electrocompetent cells (Lucigen) and grown at 30 °C for 16 h in LB with 100 µl/mL carbenicillin. Cells were harvested by centrifugation and plasmid DNA was prepared. To confirm library representation and distribution, inserts were amplified from plasmid library in a reaction of 25 µl 2x NEBnext PCR master mix, 1 µl plasmid library (10 ng/µl), 0.25 µl forward and 0.25 µl reverse primer (a 1:1:1:1:1:1 mix of primers Fw_sensor_P5_S0-Fw_sensor_P5_S5 and Rv_sensor_P7_S0-Rv_sensor_P7_S5 with various stagger lengths to create sequence diversity in the samples; Table S16) at a final concentration of 0.5 uM, and 23.5 µl water. PCR cycling conditions were: (1) 98 °C for 30 s; (2) 98 °C for 10 s; (3) 65 °C for 30 s; (4) 72 °C for 20 s; (5) go to (2) x 16; (6) 72 °C for 3 min; (7) 10 °C hold. A second PCR reaction of 25 µl NEBnext PCR master mix, 10 ng PCR1 product, 2.5 µl forward and reverse primer (Table S16) in a total of 50 µl introduced i5 and i7 multiplexing barcodes using the following PCR cycling conditions: (1) 98 °C for 30 s; (2) 98 °C for 10 s; (3) 65 °C for 75 s; (4) go to (2) x 7; (6) 72 °C for 5 min; (7) 10 °C hold. Amplicons were first gel-purified and then re-purified by PCR-purification. Samples were sequenced on a NextSeq2000 (Illumina) with a 25% spike-in of PhiX. Fastq files were generated using bcl2fastq v.2.19.1 and reads were trimmed using cutadapt 1.18, removing a variable number of basepairs at the start and end depending on the size of the primer stagger. MAGeCK^54^ 0.5.8 was used to assign the trimmed reads to the gRNAs in the library and create the gRNA count matrices.

### Virus production and competitive growth for sensor and rhAmpSeq screens

Lentiviral libraries were produced as described above and the screens were conducted as described above as a triplicate (three separate transductions) at low multiplicity of infection (MOI ∼0.3) and high coverage (>1000X) with the following alterations: A) 24 hours after transduction, transduced HAP1 cells were selected by splitting into medium with 1 μg/mL puromycin. B) The screen was terminated at day 18. Additionally, the *sensor screen* was performed in hTERT RPE1 p53-/cells (a kind gift from D. Durocher) with the following alterations compared to HAP1 cells: A) Cells were cultured in DMEM GlutaMax (Gibco) + 10% FBS (Gibco) + 100 U/mL penicillin–streptomycin (Gibco) and 0.25% trypsin-EDTA (Gibco) was used for trypsinization. B) Transduced cells were selected by splitting into 15 μg/mL puromycin and incubation for 24 hours, followed by an additional split into 15 μg/mL puromycin and incubation for 24 hours. For sensor libraries, a ∼255 bp plasmid library fragments containing the sgRNA sequences including the sensor were amplified from the genomic DNA by PCR using NEBNext Ultra II Q5 Master Mix with a 1:1:1:1:1:1 mix of primers Fw_sensor_P5_S0-Fw_sensor_P5_S5 and Rv_sensor_P7_S0-Rv_sensor_P7_S5 with various stagger lengths to create sequence diversity in the samples (Table S16). A second PCR reaction introduced i5 and i7 multiplexing barcodes (Table S16), whereafter ∼330 bp PCR products were gel-purifed and sequenced (paired end, Read 1: 190 cycles ; read 2: 75 cycles) on Illumina NextSeq2000 with a 25% spike-in of PhiX. Fastq files were generated using bcl2fastq v.2.19.1 and reads were trimmed using cutadapt 1.18, removing a variable number of basepairs at the start and end depending on the size of the primer stagger. MAGeCK^54^ 0.5.8 was used to assign the trimmed reads to the gRNAs in the libraries and create the gRNA count matrix.

### rhAmpSeq pooled amplicon sequencing

Genomic DNA from the *rhAmpSeq* screen samples were subjected to pooled amplification of the 25 genomic loci targeted by the gRNAs (Table S17) using the rhAmpSeq CRISPR analysis system (IDT) following the instructions of the manufacturer with a custom 25 primer pair rhAmpSeq CRISPR panel. Briefly, the amplification reaction consisted of 5 µl 4x rhAmpSeq CRISPR Library mix 1 (IDT), 11 µl genomic DNA (4.55 ng/µl) and 2 µl each of forward and reverse primer mixes at a concentration of 250 nM. PCR cycling conditions were: (1) 95 °C for 10 min; (2) 95 °C for 15 s; (3) 61 °C for 8 min; (4) go to (2) x 13; (5) 99.5 °C for 15 min; (6) 4 °C hold. The PCR products were diluted 1:20 in ultrapure water and immediately used as template in a second PCR reaction that introduced i5 and i7 multiplexing barcodes. The amplification reaction consisted of 5 µl 4x rhAmpSeq CRISPR Library mix 2 (IDT), 11 µl diluted PCR1 product and 2 µl each of i5 and i7 indexing primers in a unique combination at a concentration of 1 µM (Table S16). PCR cycling conditions were: (1) 95 °C for 3 min; (2) 95 °C for 15 s; (3) 60 °C for 30 s; (4) 72 °C for 30 s; (5) go to (2) x 23; (6) 72 °C for 1 min; (6) 4 °C hold. The indexed PCR reactions were pooled and purified by adding 100 µl Agencourt® AMPure® XP Beads to 100 µl pooled PCR reaction, followed by incubation for 10 min at RT and placing in a magnetic separation rack for 5 min. The beads were washed twice in 80% ethanol and dried, and the DNA eluted in 22 µl IDTE pH 8.0 by vortexing. The tube was placed in the magnetic separation rack, and the eluate was collected without beads and sequenced on Illumina NextSeq2000 (paired end, Read 1: 161 cycles ; read 2: 161 cycles) with a 25% spike-in of PhiX. Fastq files were generated using bcl2fastq v.2.19.1 and Read 1 and 2 were stiched.

### Sensor and rhAmpSeq screen data processing

*Sensor screen:* For each timepoint and replicate, paired-end NGS data were generated with the gRNA sequence located in Read 1 and the corresponding sensor sequence in Read 2. gRNA regions were extracted from Read 1 using the construct flanking sequences and reads lacking a designed gRNA were removed. gRNA counts were determined and processed as described for the initial base editing screen. For each Read 1 containing a valid gRNA, the paired Read 2 was processed by extracting the sensor region using the designed flanking sequences. Each sensor was verified against the expected barcode for its gRNA, and barcode–gRNA mismatches were discarded. Sensor sequences were then trimmed to the designed region, grouped by gRNA, and converted into gRNA-centric nucleotide-level edit count tables. These nucleotide edits were mapped to protein-level edits, including splice-site and premature stop annotations where relevant, and counts for identical protein-level editing outcomes were merged. The final output included nucleotide editing frequencies, protein editing frequencies, and detailed frequencies for specific protein, splice-site, and stop edits. Analysis focused on editing data obtained at day 3. *rhAmpSeq screen, amplicon data:* The rhAmpSeq amplicon NGS data were processed largely as described for the sensor sequence in the sensor screen with the following exceptions: a single stitched read was used; reads were trimmed using unique genomic flanking sequences to recover amplicon sequences aligned to the designed constructs; and reads corresponding to unedited amplicons were removed, as they likely originated from cells transduced with a different gRNA. *rhAmpSeq screen, sensor data:* NGS data was processed using the pipeline described for the *sensor screen*. Final editing outcome frequencies for the *rhAmpSeq screen’s* sensor and amplicon data were aggregated at the protein residue level to generate position-specific editing frequency matrices for visualization as sequence logos.

### Deep sequencing of CPSF1 locus

Individual sgRNAs targeting CPSF1 were ordered as DNA oligonucleotides (TAG Copenhagen) with overhangs for Esp3I cloning: Forward oligo: 5’-CACCG[gRNA (20nt)]-3’; Reverse oligo: 5’-AAAC[reverse complement gRNA (20nt)]C-3’ (Table S16). The oligonucleotides were annealed and phosphorylated by T4 PNK (NEB) and cloned into either base editor construct pRDA_478 or pRDA_479 using golden gate cloning with Esp3I and T7 ligase. Each sgRNA was cloned into the backbone where it was predicted to give an edit. Empty pRDA_478 and pRDA_479 were used as controls. Ligated plasmids were transformed into NEB Stable Competent E. coli and grown on LB agar containing ampicillin at 30°C for 24 h, followed by plasmid preparation from single colonies. Virus particles for each sgRNA-expressing base editor construct were produced as described above. HAP1 cells were transduced as a single biological replicate with each sgRNA-expressing base editor construct, selected, and cultured for 18 days after selection (d0-d18) as described above. Genomic DNA was extracted from cell pellets harvested at d0 and d18 using the QIAamp DNA Blood Mini Kit (Qiagen). A PCR reaction was set up to amplify the genomic CPSF1 locus, where editing was predicted to occur, and to introduce Illumina TruSeq Adaptor flaps. This was done using NEBNext Ultra II Q5 Master Mix with primers: CPSF1_Fw and CPSF1_Rv (Table S16). A second PCR reaction introduced unique combinations of Illumina i5 and i7 multiplexing barcodes using NEBNext Ultra II Q5 Master Mix and Illumina TruSeq primers listed in Table S16. Gel-purified PCR products were sequenced on Illumina NextSeq2000 with a 25% spike-in of PhiX and analysed by CRISPResso2^59^ (Supplemental Data 1).

### Cloning

pcDNA5/FRT/TO plasmids encoding venus-tagged (an enhanced YFP derivate referred to as YFP for simplicity) baits (Table S16) in a wild-type (WT) or mutant (Mut) version were generated by gene synthesis, mutagenesis, and subcloning into pcDNA5/FRT/TO/venus (GeneArt, ThermoScientific). pcDNA5/FRT/TO/Venus-CPSF1 and NCAPH were cloned from in house constructs and mutated by site-directed mutagenesis PCR. WDR33 431-485 sequences were amplified by PCR and cloned into pGEX4T-1. The E.coli expression construct for CSTF3 21-549 was done by LIC cloning into an in house expression vector. All constructs were verified by sequencing.

### Affinity purification coupled to mass spectrometry (AP-MS) sample preparation

HeLa cells were transfected with plasmids encoding YFP-tagged full length protein baits (or larger fragments in the case of ANKRD17_1950-2603_ and WDR33_1-525_) (Table S16) using Lipofectamine 2000 24 or 48 hours prior to harvesting. The bait proteins were either wild-type (WT) or motif mutant (Mut) (Fig. S9, Table S16). Cells were lysed on ice in lysis buffer (50 mM Tris pH 7.4, 150 mM NaCl, 1 mM EDTA, 0.1% NP-40) supplemented with 1 mM DTT, complete protease inhibitor cocktail (Roche), and 4U/mL RNAse, and sonicated for 10 min (30’’ON/30’’OFF, max) using Bioruptor Plus (Diagenode). Lysates were cleared by centrifugation at 20,000 g at 4°C for 30 minutes, and YFP-tagged baits were affinity purified using GFP-trap beads (ChromoTek). Purified protein complexes were washed twice in lysis buffer and once in PBS. Partial on-bead digestion was performed with buffer (2 M urea; 2 mM DTT; 20 µg/ml Trypsin; 50 mM Tris, pH 7.5) incubated at 37°C for 30 min at 1,400 rpm. Supernatants were transferred to new tubes, alkylated with 25 mM chloroacetamide (CAA) and further digested overnight at RT and 1,000 rpm. Digestion was terminated with 1% trifluoroacetic acid (TFA). Peptides were desalted and purified using styrenedivinylbenzene-reversed phase sulfonate (SDB-RPS) StageTips prepared in 0.2% TFA. Peptides were washed and eluted with an elution buffer (80% acetonitrile (ACN); 1% ammonia) prior to vacuum-drying. Dried peptides were reconstituted in 2% ACN and 0.1% TFA. Technical quadruplicates were performed.

### Liquid chromatography-mass spectrometry (LC-MS) analysis

All samples were loaded onto Evotips Pure and measured with a data-independent acquisition (DIA) method. 200 ng of peptides were loaded onto Evotips following the manufacturer’s instructions, and then analyzed with an Evosep One LC system (Evosep Biosystems) coupled online to an Orbitrap mass spectrometer (Orbitrap Astral or Orbitrap Exploris 480, Thermo Fisher Scientific)^60–62^. Eluted peptides were separated on a 8-cm-long PepSep column (150 µm inner diameter packed with 1.5 μm of Reprosil-Pur C18 beads (Dr Maisch)) in a standard gradient method with a stainless emitter (30 µm inner diameter). The mobile phases were 0.1% formic acid in liquid chromatography (LC)–MS-grade water (buffer A) and 0.1% formic acid in acetonitrile (buffer B). Data was acquired in DIA mode.

For the samples acquired in the Orbitrap Exploris 480, peptides were eluted online from the EvoTips using an Evosep One LC system and analysed at 30 samples per day (44-min gradient). The mass spectrometer was operated in positive mode using the DIA mode. Full scan precursor spectra (350–1,000 m/z) were recorded in profile mode using a resolution of 120,000, with a normalised automatic gain control target (AGC) of 300% and a maximum injection time of 45 ms. Fragment spectra were then recorded in profile mode, fragmenting 49 consecutive 13.3 m/z windows covering the mass range 350–1,000 m/z and using a resolution of 15,000. Isolated precursors were fragmented in the higher-energy collisional dissociation (HCD) cell using 27% normalised collision energy, a normalised AGC target of 3,000 % and a maximum injection time of 22 ms.

For the samples acquired in the Orbitrap Astral, peptides were eluted online from the EvoTips using an Evosep One LC system and analysed at 60 samples per day (21-min gradient). The Orbitrap Astral mass spectrometer was operated in positive mode using the DIA mode, at a resolution of 240,000 with a full scan range of 380-980 m/z. The normalised AGC was set to 500%. Fragment ion scans were recorded at a maximum injection time of 3.5 ms, 200 windows of 3Th scanning from 380-980 m/z. Ions were fragmented using HCD with 25% normalised collision energy.

### Mass spectrometry raw data processing and bioinformatic analysis

Raw files were analysed with directDIA workflow in Spectronaut v.19.0^63^. Default settings were used. Data filtering was set to ‘Qvalue’. ‘Cross run normalisation’ was enabled with the strategy of ‘local normalisation’ based on rows with ‘Qvalue complete’. FDR was set to 1% at both the protein and peptide precursor levels. Raw data was searched against the human proteome reference database (including 20,593 entries, Uniprot March 2023). Data was filtered for 75% valid values (in at least 3 out 4 replicates) across WT or Mutant samples (protein groups with >25% missing values were excluded from downstream statistical analysis). Protein intensities were then log2 transformed for downstream statistical and bioinformatics analysis. For each protein, missing values were sampled from a normal distribution with the following parameters: mean = median of all quantified values across all proteins in each sample minus 1.8 times the standard deviation, standard deviation = the standard deviation of all quantified values across all proteins in each sample multiplied by 0.3. For ZNHIT-IP, we did extra z-score normalization to reduce the technical variations. The significance of protein enrichment of Mutants relative to WT was determined through an unpaired two-sided *t*-test corrected by Benjamini–Hochberg at FDR of 0.1 and fold change of 1.5^64,65^. Data analysis and visualisation were done in Jupyter Notebook in the environment of Python 3.9.

### AP-WB

HeLa cells were transfected with plasmids encoding the specified YFP-tagged baits in a wild-type version or with the indicated mutations using Lipofectamine 2000 24 hours prior to harvesting. Cells were lysed on ice in lysis buffer (50 mM Tris pH 7.4, 150 mM NaCl, 1 mM EDTA, 0.1% NP-40) supplemented with 1 mM DTT and complete protease inhibitor cocktail (Roche) and sonicated for 10 min (30’’ON/30’’OFF, max) using Bioruptor Plus (Diagenode). Lysates were cleared by centrifugation at 20,000 g at 4°C for 30 minutes, and YFP-tagged baits were affinity purified using GFP-trap beads (ChromoTek). Precipitated protein complexes were washed thrice in lysis buffer and eluted in 2X LDS sample buffer before SDS-PAGE and Western blot analysis with the indicated antibodies (CSTF3: HPA040168 (Sigma), CNOT1: 14276-1-AP (proteintech), YFP: GFP in house made antibody to full length GFP). Uncropped WBs for panel 4e and 5i are shown in Fig. S11.

### Protein production

CSTF3_21-549_ wild-type and GST-WDR33_431-485_ wild-type and mutant were expressed in BL21(DE3) cells overnight at 18 degrees and cells were harvested by centrifugation. For CSTF3 purification cells were resuspended in Buffer L (50 mM NaP pH 7.5; 300 mM NaCl; 10 mM imidazole; 10% glycerol; 0.5 mM TCEP; 1xComplete EDTA-free tablets (Roche)) and lysed. Following clarification by centrifugation, lysate was loaded on a 5 ml HiTrap column and His-CSTF3_21-549_ eluted with an imidazole gradient. The peak fractions were pooled, diluted with 50 mM NaP pH 7.5 to 50 mM NaCl and loaded on a SP HP HiTrap column and proteins eluted with a 50mM-1M NaCl salt gradient. Peak fractions were pooled.

For purification of GST-WDR33_431-485_, cell pellets were resuspended in 25 mL lysis buffer (50 mM Tris-HCl pH 7.4, 300 mM NaCl, 10% glycerol, 5 mM β-mercaptoethanol) and sonicated for 10 minutes on ice. Lysates were cleared in a fixed angle rotor for 40 min 145000 rpm and supernatants were incubated with 350 µL pre-equilibrated Glutathione Sepharose Fast Flow (GE healthcare) for 1h. Beads were washed 3 times with 30 mL lysis buffer and used directly for binding experiments.

### GST-WDR33 pull down assay

10 µL GST-WDR3_431-485_ wild-type or mutant beads in 200 µL buffer B (150 mM NaCl, 50 mM Tris pH 7.4 1 mM DTT) were incubated with 20 µg CTSF3 for 1h at 4C with rotation. Beads were washed 2 times with buffer B and eluted in 25 µL 2x LDS sample buffer (Thermo Fisher Scientific) with 10 mM DTT. For control conditions unbound glutathione beads were used. Data is obtained from three replicates.

### ITC

Peptides were purchased from Peptide 2.0 Inc (Chantilly. VA, USA). The purity obtained in the synthesis was 95–98% as determined by high performance liquid chromatography (HPLC) and subsequent analysis by mass spectrometry. Prior to ITC experiments both CSTF3*^21–549^* and the peptides were extensively dialyzed against 50 mM sodium phosphate pH 7.5, 150 mM NaCl, 0.5 mM TCEP. CSTF3*^21-549^*and peptide concentrations were determined using a spectrometer by measuring the absorbance at 280 nm and applying values for the extinction coefficients computed from the corresponding sequences by the ProtParam program (http://web.expasy.org/protparam/). Peptides at approximately 500 μM concentration were loaded into the syringe and titrated into the calorimetric cell containing CSTF3*^21-549^* at ∼ 40 μM (monomer concentration). The reference cell was filled with distilled water. In all assays, the titration sequence consisted of a single 0.4 μl injection followed by 19 injections, 2 μl each, with 150 s spacing between injections to ensure that the thermal power returns to the baseline before the next injection. The stirring speed was 750 rpm. Control experiments with peptides injected in the sample cell filled with buffer were carried out under the same experimental conditions. These control experiments showed negligible heats of dilution. Experiments were caried out as a single replicate. The heats per injection normalised per mole of the injectant *versus* the molar ratio [Peptide]/[CSTF3*^21-549^*] were fitted to a single-site model. Data were analysed with MicroCal PEAQ-ITC (version 1.1.0.1262) analysis software (Malvern Instruments Ltd.).

### AlphaFold predictions

AlphaFold models were generated by submitting full length protein sequences of CPSF1, WDR33, CPSF4, CSTF3 and PAS-RNA (inspired by the human Pre-mRNA 3’-End Processing Machinery structure PDB:6URO) to AlphaFold3^66^ (https://alphafoldserver.com) and was ran using standard parameters without templates. Overall predicted template modelling (PTM) and interface predicted template modelling (iPTM) scores were 0.67 and 0.74, respectively. The 5 predictions were further inspected for confidence in ChimeraX^67^ by aligning the predictions to the PDB:6URO structure^42^ using the *matchmaker* function. Confidence of the CPSF1_742-749_ CSTF3 interaction was assessed by aligning the models to the FIP1L1_1-35_-CSTF3 crystal structure (PDB: 7ZY4)^38^, confirming the same CPSF1-CSTF3 interaction mode for all 5 predictions, which for the motifs was highly similar to that of FIP1L1. For the WDR33_441-445_ motif, confidence of the predictions was confirmed given that all 5 predictions modelled the IPGMG_441-445_ in the same binding pocket of CSTF3 with average atom pLDDT of IPGMG_441-445_ ranging from 64.44-65.57.

## Data availability

The proteomic raw data is available at PRIDE ID PXD055750

Username

Password: lyWJc1nnFmbq

All screen sequencing data is available as gRNA count files in supplementary tables.

## Code availability

Code will be made available upon request.

## Bibliography

1. Tompa, P., Davey, N.E., Gibson, T.J. & Babu, M.M. A million peptide motifs for the molecular biologist. Mol Cell 55, 161–9 (2014).

2. Van Roey, K. et al. Short linear motifs: ubiquitous and functionally diverse protein interaction modules directing cell regulation. Chem Rev 114, 6733–78 (2014).

3. Davey, N.E., Simonetti, L. & Ivarsson, Y. The next wave of interactomics: Mapping the SLiM-based interactions of the intrinsically disordered proteome. Curr Opin Struct Biol 80, 102593 (2023).

4. Davey, N.E. et al. Attributes of short linear motifs. Mol Biosyst 8, 268–81 (2012).

5. Kumar, M. et al. ELM-the Eukaryotic Linear Motif resource-2024 update. Nucleic Acids Res 52, D442–d455 (2024).

6. Holehouse, A.S. & Kragelund, B.B. The molecular basis for cellular function of intrinsically disordered protein regions. Nat Rev Mol Cell Biol 25, 187–211 (2024).

7. Teilum, K., Olsen, J.G. & Kragelund, B.B. On the specificity of protein-protein interactions in the context of disorder. Biochem J 478, 2035–2050 (2021).

8. Davey, N.E., Cyert, M.S. & Moses, A.M. Short linear motifs ex nihilo evolution of protein regulation. Cell Commun Signal 13, 43 (2015).

9. Kliche, J. et al. Proteome-scale characterisation of motif-based interactome rewiring by disease mutations. Mol Syst Biol 20, 1025–1048 (2024).

10. Mészáros, B., Kumar, M., Gibson, T.J., Uyar, B. & Dosztányi, Z. Degrons in cancer. Sci Signal 10(2017).

11. Roy, J., Li, H., Hogan, P.G. & Cyert, M.S. A conserved docking site modulates substrate affinity for calcineurin, signaling output, and in vivo function. Mol Cell 25, 889–901 (2007).

12. Lue, N.Z. & Liau, B.B. Base editor screens for in situ mutational scanning at scale. Mol Cell 83, 2167–2187 (2023).

13. Richter, M.F., et al. Phage-assisted evolution of an adenine base editor with improved Cas domain compatibility and activity. Nat Biotechnol 38, 883–891 (2020).

14. Koblan, L.W., et al. Improving cytidine and adenine base editors by expression optimization and ancestral reconstruction. Nat Biotechnol 36, 843–846 (2018).

15. Walton, R.T., Christie, K.A., Whittaker, M.N. & Kleinstiver, B.P. Unconstrained genome targeting with near-PAMless engineered CRISPR-Cas9 variants. Science 368, 290–296 (2020).

16. Sangree, A.K., et al. Benchmarking of SpCas9 variants enables deeper base editor screens of BRCA1 and BCL2. Nat Commun 13, 1318 (2022).

17. Hanna, R.E., et al. Massively parallel assessment of human variants with base editor screens. Cell 184, 1064–1080.e20 (2021).

18. Ritchie, M.E. et al. limma powers differential expression analyses for RNA-sequencing and microarray studies. Nucleic Acids Res 43, e47 (2015).

19. Tunyasuvunakool, K. et al. Highly accurate protein structure prediction for the human proteome. Nature 596, 590–596 (2021).

20. Blomen, V.A. et al. Gene essentiality and synthetic lethality in haploid human cells. Science 350, 1092–6 (2015).

21. Safaee, N. et al. Interdomain allostery promotes assembly of the poly(A) mRNA complex with PABP and eIF4G. Mol Cell 48, 375–86 (2012).

22. Clute, P. & Pines, J. Temporal and spatial control of cyclin B1 destruction in metaphase. Nat Cell Biol 1, 82–7 (1999).

23. Lee, E.F. et al. The functional differences between pro-survival and proapoptotic B cell lymphoma 2 (Bcl-2) proteins depend on structural differences in their Bcl-2 homology 3 (BH3) domains. J Biol Chem 289, 36001–17 (2014).

24. Coutandin, D. et al. Conformational stability and activity of p73 require a second helix in the tetramerization domain. Cell Death Differ 16, 1582–9 (2009).

25. Czabotar, P.E. et al. Mutation to Bax beyond the BH3 domain disrupts interactions with pro-survival proteins and promotes apoptosis. J Biol Chem 286, 7123–31 (2011).

26. Suzuki, M., Youle, R.J. & Tjandra, N. Structure of Bax: coregulation of dimer formation and intracellular localization. Cell 103, 645–54 (2000).

27. Meldal, B.H.M., et al. Complex Portal 2018: extended content and enhanced visualization tools for macromolecular complexes. Nucleic Acids Res 47, D550–d558 (2019).

28. Sánchez-Rivera, F.J. et al. Base editing sensor libraries for high-throughput engineering and functional analysis of cancer-associated single nucleotide variants. Nature Biotechnology 40, 862–873 (2022).

29. Kim, Y., et al. High-throughput functional evaluation of human cancer-associated mutations using base editors. Nat Biotechnol 40, 874–884 (2022).

30. Song, M., et al. Sequence-specific prediction of the efficiencies of adenine and cytosine base editors. Nat Biotechnol 38, 1037–1043 (2020).

31. Ryu, J. et al. Joint genotypic and phenotypic outcome modeling improves base editing variant effect quantification. Nat Genet 56, 925–937 (2024).

32. Sellés-Baiget, S. et al. Catalytic and noncatalytic functions of DNA polymerase κ in translesion DNA synthesis. Nat Struct Mol Biol 32, 300–314 (2025).

33. Arafeh, R., Shibue, T., Dempster, J.M., Hahn, W.C. & Vazquez, F. The present and future of the Cancer Dependency Map. Nat Rev Cancer 25, 59–73 (2025).

34. Chaikovsky, A.C. et al. The AMBRA1 E3 ligase adaptor regulates the stability of cyclin D. Nature 592, 794–798 (2021).

35. Maiani, E. et al. AMBRA1 regulates cyclin D to guard S-phase entry and genomic integrity. Nature 592, 799–803 (2021).

36. Simoneschi, D. et al. CRL4(AMBRA1) is a master regulator of D-type cyclins. Nature 592, 789–793 (2021).

37. Lei, Q.Y. et al. TAZ promotes cell proliferation and epithelial-mesenchymal transition and is inhibited by the hippo pathway. Mol Cell Biol 28, 2426–36 (2008).

38. Muckenfuss, L.M., Migenda Herranz, A.C., Boneberg, F.M., Clerici, M. & Jinek, M. Fip1 is a multivalent interaction scaffold for processing factors in human mRNA 3’ end biogenesis. Elife 11 (2022).

39. Yamazaki, Y. et al. Goliath family E3 ligases regulate the recycling endosome pathway via VAMP3 ubiquitylation. Embo j 32, 524–37 (2013).

40. von Moeller, H. et al. Structural and biochemical studies of SLIP1-SLBP identify DBP5 and eIF3g as SLIP1-binding proteins. Nucleic Acids Res 41, 7960–71 (2013).

41. Udupa, A., Kotha, S.R. & Staller, M.V. Commonly asked questions about transcriptional activation domains. Curr Opin Struct Biol 84, 102732 (2024).

42. Zhang, Y., Sun, Y., Shi, Y., Walz, T. & Tong, L. Structural Insights into the Human Pre-mRNA 3’-End Processing Machinery. Mol Cell 77, 800–809.e6 (2020).

43. Hernández, M.V., Wehrendt, D.P. & Arregui, C.O. The protein tyrosine phosphatase PTP1B is required for efficient delivery of N-cadherin to the cell surface. Mol Biol Cell 21, 1387–97 (2010).

44. Benz, C. et al. Proteome-scale mapping of binding sites in the unstructured regions of the human proteome. Mol Syst Biol 18, e10584 (2022).

45. Castello, A. et al. Insights into RNA Biology from an Atlas of Mammalian mRNA-Binding Proteins. Cell 149, 1393–1406 (2012).

46. Chopra, M. et al. Heterozygous ANKRD17 loss-of-function variants cause a syndrome with intellectual disability, speech delay, and dysmorphism. Am J Hum Genet 108, 1138–1150 (2021).

47. Boland, A. et al. Structure and assembly of the NOT module of the human CCR4NOT complex. Nat Struct Mol Biol 20, 1289–97 (2013).

48. Youn, J.Y. et al. High-Density Proximity Mapping Reveals the Subcellular Organization of mRNA-Associated Granules and Bodies. Mol Cell 69, 517–532.e11 (2018).

49. Corbi-Verge, C. & Kim, P.M. Motif mediated protein-protein interactions as drug targets. Cell Commun Signal 14, 8 (2016).

50. Monzon, A.M., Bonato, P., Necci, M., Tosatto, S.C.E. & Piovesan, D. FLIPPER: Predicting and Characterizing Linear Interacting Peptides in the Protein Data Bank. J Mol Biol 433, 166900 (2021).

51. Davey, N.E. et al. SLiMPrints: conservation-based discovery of functional motif fingerprints in intrinsically disordered protein regions. Nucleic Acids Res 40, 10628–41 (2012).

52. Dosztányi, Z., Csizmók, V., Tompa, P. & Simon, I. The pairwise energy content estimated from amino acid composition discriminates between folded and intrinsically unstructured proteins. J Mol Biol 347, 827–39 (2005).

53. UniProt Consortium, T. UniProt: the universal protein knowledgebase. Nucleic Acids Res 46, 2699 (2018).

54. Li, W. et al. MAGeCK enables robust identification of essential genes from genome-scale CRISPR/Cas9 knockout screens. Genome Biol 15, 554 (2014).

55. Kotb, H.M. & Davey, N.E. xProtCAS: A Toolkit for Extracting Conserved Accessible Surfaces from Protein Structures. Biomolecules 13 (2023).

56. Mi, H. et al. Protocol Update for large-scale genome and gene function analysis with the PANTHER classification system (v.14.0). Nat Protoc 14, 703–721 (2019).

57. Tsitsa, I., Krystkowiak, I. & Davey, N.E. CompariPSSM: a PSSM-PSSM comparison tool for motif-binding determinant analysis. Bioinformatics 40(2024).

58. Davey, N.E., Edwards, R.J. & Shields, D.C. The SLiMDisc server: short, linear motif discovery in proteins. Nucleic Acids Res 35, W455–9 (2007).

59. Clement, K. et al. CRISPResso2 provides accurate and rapid genome editing sequence analysis. Nat Biotechnol 37, 224–226 (2019).

60. Bache, N. et al. A Novel LC System Embeds Analytes in Pre-formed Gradients for Rapid, Ultra-robust Proteomics. Mol Cell Proteomics 17, 2284–2296 (2018).

61. Heil, L.R. et al. Evaluating the Performance of the Astral Mass Analyzer for Quantitative Proteomics Using Data-Independent Acquisition. J Proteome Res 22, 3290–3300 (2023).

62. Guzman, U.H. et al. Ultra-fast label-free quantification and comprehensive proteome coverage with narrow-window data-independent acquisition. Nat Biotechnol 42, 1855–1866 (2024).

63. Bruderer, R. et al. Extending the limits of quantitative proteome profiling with data-independent acquisition and application to acetaminophen-treated three-dimensional liver microtissues. Mol Cell Proteomics 14, 1400–10 (2015).

64. Lin, M.H. et al. Benchmarking differential expression, imputation and quantification methods for proteomics data. Brief Bioinform 23(2022).

65. Dowell, J.A., Wright, L.J., Armstrong, E.A. & Denu, J.M. Benchmarking Quantitative Performance in Label-Free Proteomics. ACS Omega 6, 2494–2504 (2021).

66. Abramson, J. et al. Accurate structure prediction of biomolecular interactions with AlphaFold 3. Nature 630, 493–500 (2024).

67. Goddard, T.D. et al. UCSF ChimeraX: Meeting modern challenges in visualization and analysis. Protein Sci 27, 14–25 (2018).

