## Supplementary figures for "A proteome-wide dependency map of protein interaction motifs"

### Supplemental material

**Supplemental figures:**

Fig. S1: Relating to Fig. 1. Benchmarking of base editing screens.

Fig. S2: Relating to Fig. 1. Base editing of the CPSF1 LYG locus.

Fig. S3: Relating to Fig. 2. Interface of the web resource to explore the data.

Fig. S4: Relating to Fig. 2. PANTHER analysis of Reactome pathway enrichment and complex analysis.

Fig. S5: Relating to Fig. 3. Sensor screen validates gRNA performance and editing across HAP1 and RPE1.

Fig. S6: Relating to Fig. 3. rhAmpSeq screen validates genomic locus editing.

Fig. S7: Relating to Fig. 4. A novel SLiM in CSTF-CPSF cleavage and polyadenylation complex is required for complex formation.

Fig. S8: Relating to Fig. 4. Mutations in CPSF1 LYG motif cause loss of binding to CSTF3.

Fig. S9: Relating to Fig. 5. Mutation of predicted SLiMs disrupt protein interactions.

Fig. S10: Relating to Fig. 5. Figure 10. SLiMs in WDR33 directly binds CSTF3.

Fig. S11: Relating to Fig. 4 and 5. Uncropped Western blots.

**Supplemental data:**

Supplemental data: Base editing of the CPSF1 LYG locus (CRISPResso2 analysis).

**Supplemental tables:**

Table S1: Target SLiM-containing regions.

Table S2: gRNA design.

Table S3: gRNA level BE screen data.

Table S4: CRISPResso2 analysis of CPSF1 LYG locus (above 10% edits).

Table S5: Region level BE screen data.

Table S6: Pathway enrichment analysis.

Table S7: Protein complex analysis.

Table S8: Sensor and rhAmpSeq gRNA libraries

Table S9: gRNA level sensor screen data.

Table S10: Region level sensor screen data.

Table S11: rhAmpSeq and sensor screen data

Table S12: AP-MS data.

Table S13: Isothermal titration calorimetry data.

Table S14: Specificity determinant analysis.

Table S15: Disease mutations overlapping with essential motifs.

Table S16: Primers and plasmids.

Table S17: rhAmpseq gRNA genomic coordinates.

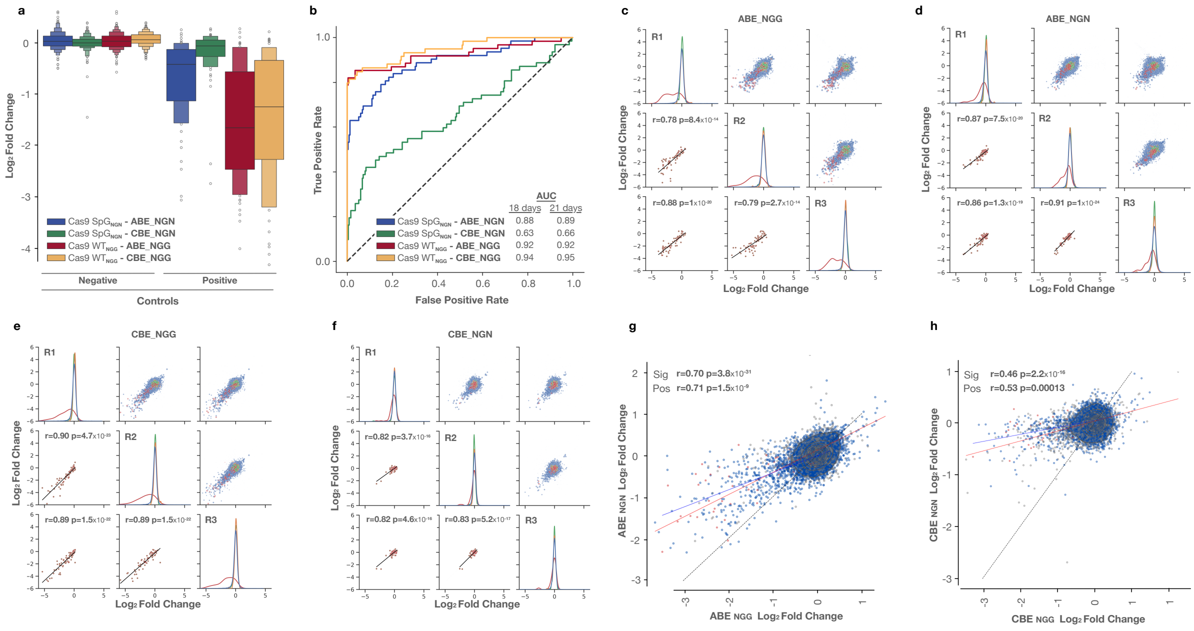


#### Supplementary figure 1: Benchmarking of base editing screens

**a**, Boxplots comparing the gRNA $\log_2$ fold changes of the positive and negative controls of the four screens for day 18 versus day 0. **b**, Receiver Operating Characteristic (ROC) curve analysis comparing the gRNA $\log_2$ fold changes of positive and negative controls of the four screens for day 21 vs day 0, inlay of the area under the curve (AUC) for day 18 and day 21. **c-f**, Correlation of the three replicates (R1-R3) from $ABE_{NGG}$ (**c**), $ABE_{NGN}$ (**d**), $CBE_{NGG}$ (**e**) and $CBE_{NGN}$ (**f**) screens. Pearson correlation coefficients (r) and p-values (p) are indicated. **g**, Comparison of the gRNA $\log_2$ fold changes for the $ABE_{NGN}$ and $ABE_{NGG}$ screens. Pearson correlation coefficients (r) and p-values (p) are indicated for significant gRNAs (sig) and positive controls (pos). **h,** As g, for the CBE screens.

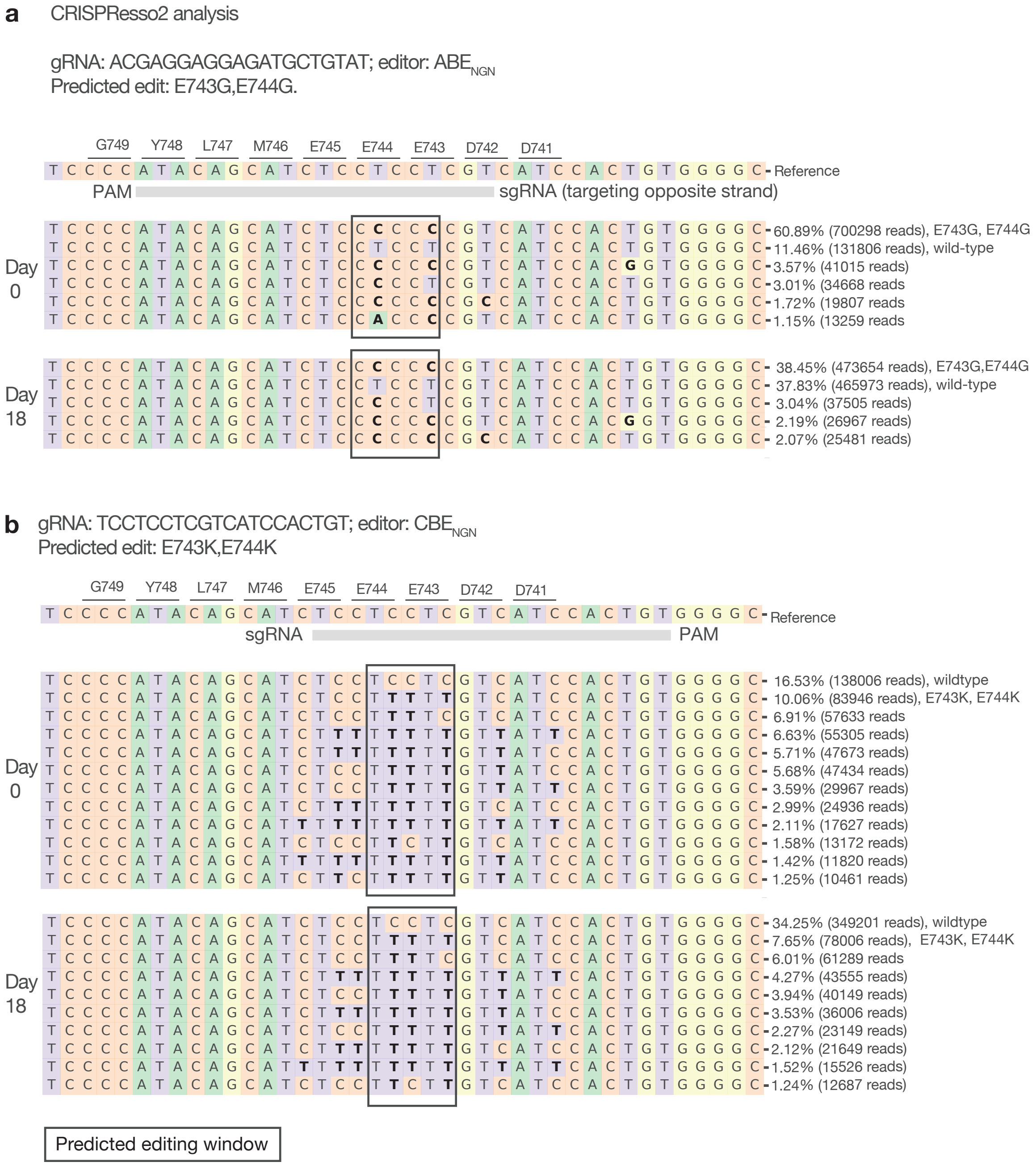


**Supplementary figure 2: Base editing of the CPSF1 LYG locus**

**a-b**, HAP1 cells transduced with the indicated single gRNAs in combination with ABE$_{NGN}$ (**a**) or CBE$_{NGN}$ (**b**) were passaged for 18 days post selection, and genomic DNA was extracted at day 0 and day 18. The LYG locus of CPSF1 was PCR amplified and subjected to NGS and subsequent CRISPResso2 analysis. Top, the reference sequence represents the wild-type locus with amino acid translation, location of the gRNA, and predicted edit indicated. In (**a**), the gRNA targets the complementary strand. Bottom, the allelic frequencies and NGS reads (above 1%) from CRISPResso2 analysis at day 0 and day 18. Translations are indicated for the most frequent mutation. The predicted editing window is boxed.

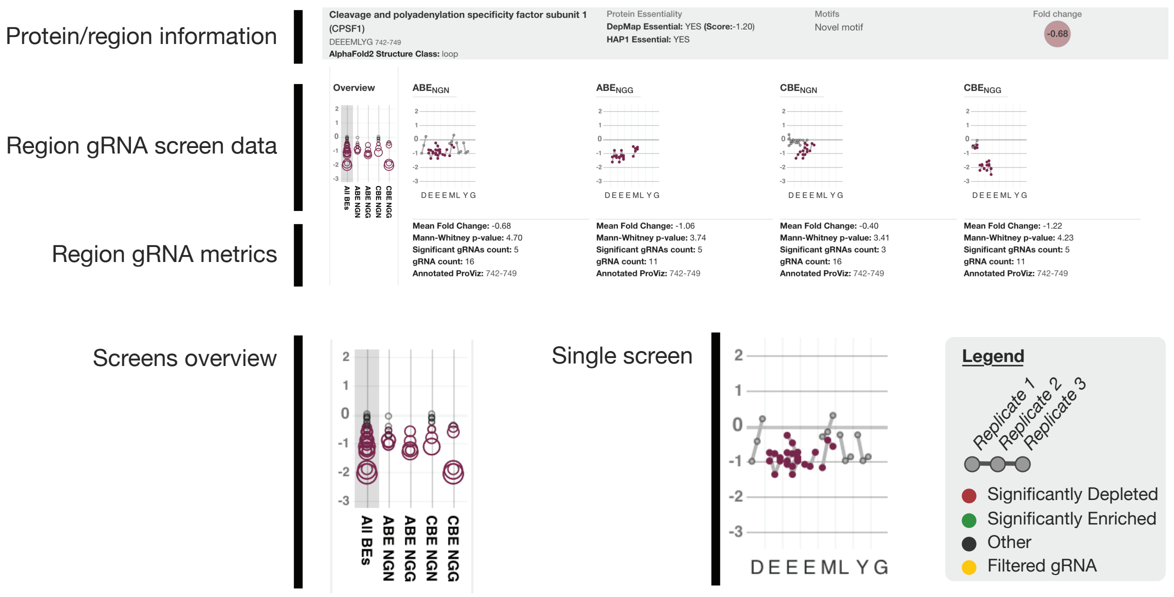


**Supplementary figure 3: Interface of the web resource to explore the data.**

Example of data that can be viewed in the web resource with CPSF1 motif as an example. Plots have y-axes indicating gRNA $\log_2$-fold changes either collected for all screens as in the 'overview' with circle size representing limma p-values (left) or for each screen in the context of the targeted amino acid residue within the presented motif (indicated on the x-axis). Connected dots indicate the same gRNA edit across the biological triplicate. In case gRNAs are predicted to target multiple residues, $\log_2$ fold changes of that gRNA are represented for each residue.

a

| FDR | Exp | Obs | Enr | Pathway |
|---|---|---|---|---|
| $6.5{\cdot}10^{-12}$ | 67.53 | 130 | 1.9 | RNA Polymerase II Transcription |
| $5.2{\cdot}10^{-12}$ | 76.01 | 141 | 1.83 | Gene expression (Transcription) |
| $6.4{\cdot}10^{-11}$ | 60.49 | 117 | 1.9 | Generic Transcription Pathway |
| $9.0{\cdot}10^{-9}$ | 41.77 | 84 | 2.01 | Cell Cycle |
| $2.6{\cdot}10^{-8}$ | 23.2 | 56 | 2.37 | Transcriptional Regulation by TP53 |
| $8.8{\cdot}10^{-7}$ | 8.48 | 27 | 3.18 | Mitotic G1 phase and G1/S transition |
| $8.0{\cdot}10^{-7}$ | 2.4 | 14 | 5.42 | Activation of BH3-only proteins |
| $9.8{\cdot}10^{-7}$ | 4.32 | 19 | 4.17 | Intrinsic Pathway for Apoptosis |
| $3.6{\cdot}10^{-6}$ | 32.48 | 64 | 1.97 | Cell Cycle, Mitotic |
| $3.7{\cdot}10^{-6}$ | 88.81 | 139 | 1.53 | Disease |
| $6.9{\cdot}10^{-6}$ | 35.36 | 67 | 1.89 | Cellular responses to stress |
| $7.1{\cdot}10^{-6}$ | 35.52 | 67 | 1.89 | Cellular responses to stimuli |
| $1.2{\cdot}10^{-5}$ | 6.88 | 22 | 3.2 | G1/S Transition |
| $1.7{\cdot}10^{-5}$ | 10.88 | 30 | 2.67 | Apoptosis |
| $3.8{\cdot}10^{-5}$ | 32 | 60 | 1.87 | Metabolism of RNA |

Apoptosis · Cell Cycle · Transcription · DNA Damage · Translation · Other

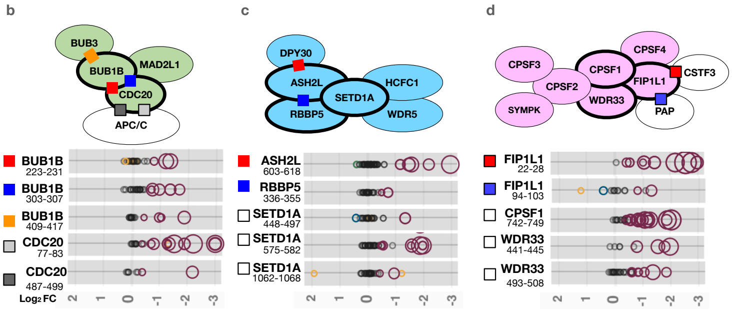


**Supplementary figure 4: Reactome pathway and complex analysis**

**a**, PANTHER analysis of Reactome pathway enrichment for genes harbouring essential SLiMs from the *reported motif set* using all the SLiM-containing genes of the *reported motif set* as the background set. FDR: false discovery rate. Exp: expected. Obs: observed. Enr: Enrichment. **b-d**, Schematic representation and gRNA $log_2$ fold changes of three protein complexes containing multiple SLiMs scoring in the screen. Colored squares indicate interaction sites of validated motifs with their binding partners while open squares indicate predicted motifs scoring in the screen. Circles in scatterplots represent a single gRNAs, with their $log_2$ fold change on the x-axis and size indicating their significance, across the 4 screens. Red, significant depletion; blue, significant enrichment; grey, stable; and yellow, filtered gRNAs.

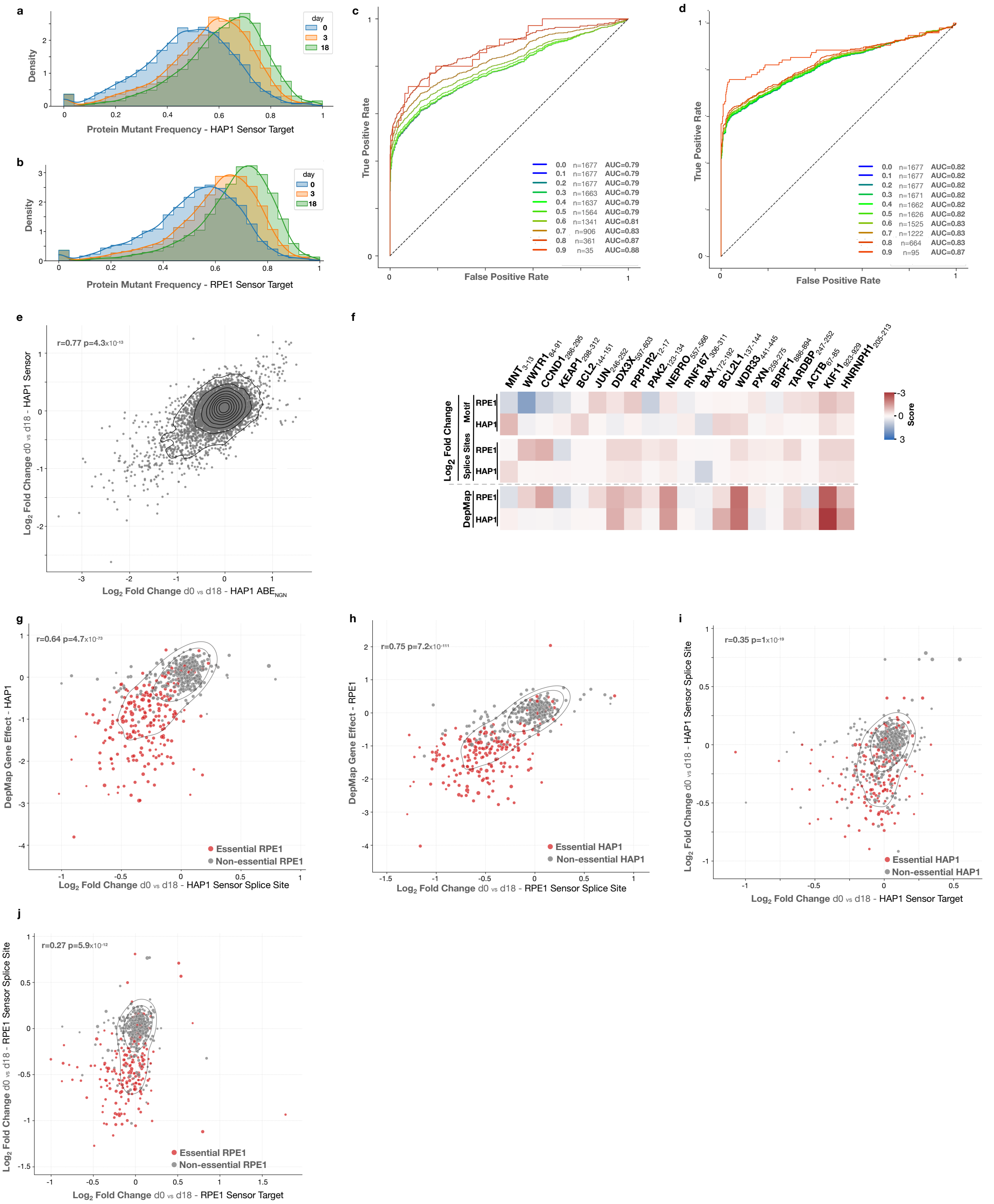
a
Density
day
0
3
18
Protein Mutant Frequency - HAP1 Sensor Target
b
Density
Protein Mutant Frequency - RPE1 Sensor Target
c
True Positive Rate
False Positive Rate
0.0 n=1677 AUC=0.79
0.1 n=1677 AUC=0.79
0.2 n=1677 AUC=0.79
0.3 n=1663 AUC=0.79
0.4 n=1637 AUC=0.79
0.5 n=1564 AUC=0.79
0.6 n=1341 AUC=0.81
0.7 n=906 AUC=0.83
0.8 n=361 AUC=0.87
0.9 n=35 AUC=0.88
d
True Positive Rate
False Positive Rate
0.0 n=1677 AUC=0.82
0.1 n=1677 AUC=0.82
0.2 n=1677 AUC=0.82
0.3 n=1671 AUC=0.82
0.4 n=1662 AUC=0.82
0.5 n=1626 AUC=0.82
0.6 n=1525 AUC=0.83
0.7 n=1222 AUC=0.83
0.8 n=664 AUC=0.83
0.9 n=95 AUC=0.87
e
r=0.77 p=4.3x10-13
Log2 Fold Change d0 vs d18 - HAP1 Sensor
Log2 Fold Change d0 vs d18 - HAP1 ABENGN
f
MNT WWTR1 CCND1 KEAP1 BCL2 JUN DDX3X PPP1R2 PAK2 NEPRO RNF167 BAX BCL2L1 WDR33 PXN BRPF1 TARDBP ACTB KIF11 HNRNPH1
Log2 Fold Change
Motif
Splice Sites
DepMap
RPE1
HAP1
Score
g
r=0.64 p=4.7x10-73
DepMap Gene Effect - HAP1
Essential RPE1
Non-essential RPE1
Log2 Fold Change d0 vs d18 - HAP1 Sensor Splice Site
h
r=0.75 p=7.2x10-111
DepMap Gene Effect - RPE1
Essential HAP1
Non-essential HAP1
Log2 Fold Change d0 vs d18 - RPE1 Sensor Splice Site
i
r=0.35 p=1x10-19
Log2 Fold Change d0 vs d18 - HAP1 Sensor Splice Site
Essential HAP1
Non-essential HAP1
Log2 Fold Change d0 vs d18 - HAP1 Sensor Target
j
r=0.27 p=5.9x10-12
Log2 Fold Change d0 vs d18 - RPE1 Sensor Splice Site
Essential RPE1
Non-essential RPE1
Log2 Fold Change d0 vs d18 - RPE1 Sensor Target

**Supplementary figure 5: Sensor screen validates gRNA performance and editing across HAP1 and RPE1**

**a**, Non-synonymous sensor editing frequency at day 0, 3, and 18 in HAP1. **b**, As a, for RPE1. **c**, Receiver operating characteristic (ROC) curves based on the gRNA fold change for the positive and negative controls in HAP1, indicating true positive rate (y-axis) vs false positive rate (x-axis) at different sensor editing frequency cut-offs (0.0 = above 0% editing, 0.9 = above 90 % editing). **d**, As c, for RPE1. **e,** Scatterplot of gRNA $log_2$ fold changes for the HAP1 $ABE_{NGN}$ screen (d18 vs d0, x-axis) versus the HAP1 sensor screen (d18 vs d0, y-axis). Pearson correlation coefficient (r) and p-value (p) are indicated. **f,** Heatmap of SLiMs with the most different $log_2$ fold changes for HAP1 vs RPE1. Gene Effect from DepMap and splice site and motif $log_2$ fold-changes from the sensor screen are indicated. Scalebar represents Gene Effect for DepMap and $log_2$ fold-changes for screen data, blue being enriched and red being depleted. **g,** Scatterplot of splice site mean $log_2$ fold changes from HAP1 (d18 vs d0, x-axis) versus HAP1 DepMap Gene Effect. Red dots indicate RPE1 essential genes from DepMap. **h**, As f, for RPE1 with red dots indicating HAP1 essential genes from DepMap. **i**, Scatterplot comparing $log_2$ fold changes of SLiMs ('target'; d18 vs d0, x-axis) with $log_2$ fold changes of their respective splice sites (d18 vs d0, y-axis) from HAP1. Red dots indicate HAP1 essential genes from DepMap . **j**, As h, for RPE1.

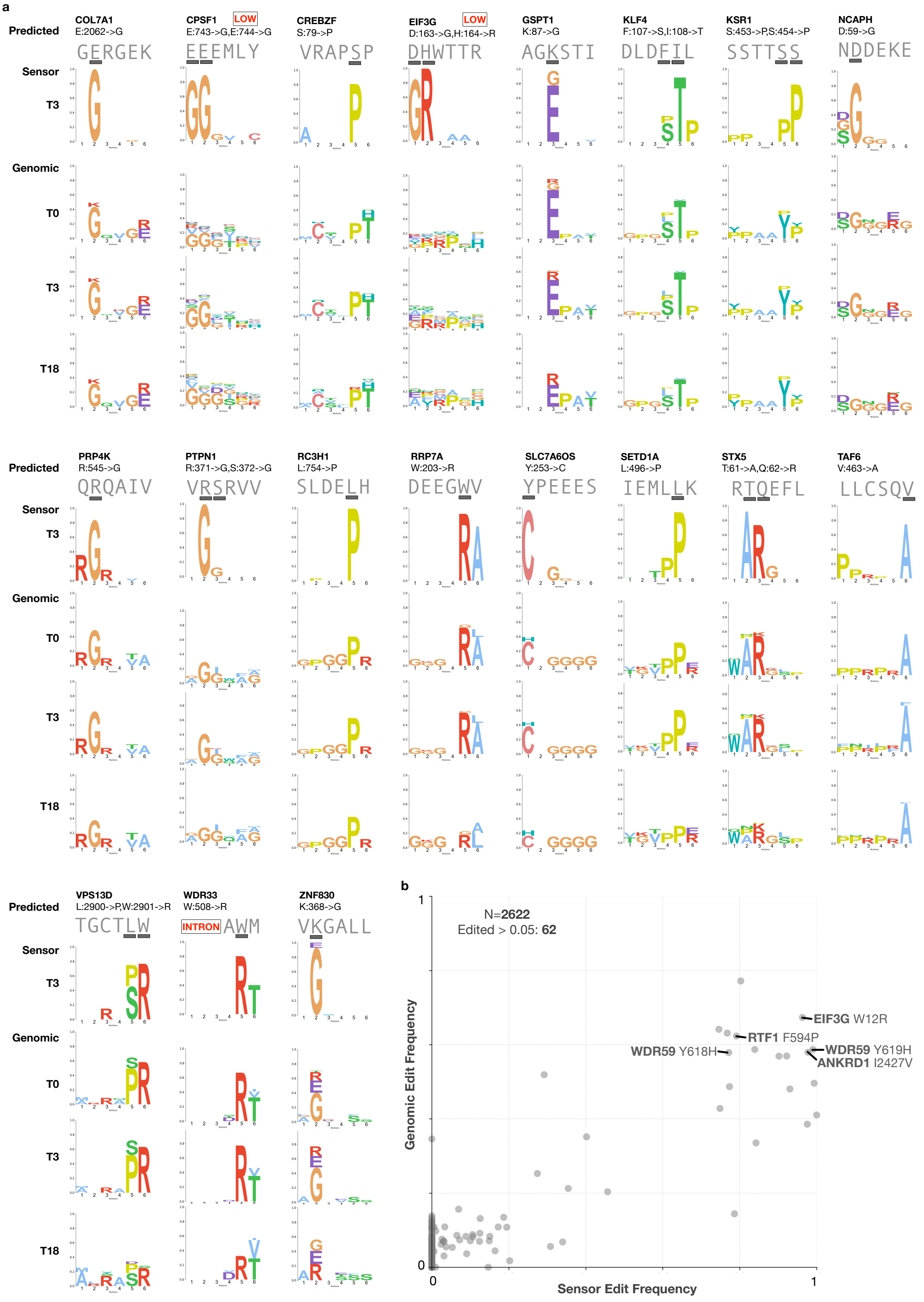
a
Predicted
Sensor
T3
Genomic
T0
T3
T18
COL7A1
E:2062->G
GERGEK
CPSF1 LOW
E:743->G,E:744->G
EEEMLY
CREBZF
S:79->P
VRAPSP
EIF3G LOW
D:163->G,H:164->R
DHWTTR
GSPT1
K:87->G
AGKSTI
KLF4
F:107->S,I:108->T
DLDFIL
KSR1
S:453->P,S:454->P
SSTTSS
NCAPH
D:59->G
NDDEKE
PRP4K
R:545->G
QRQAIV
PTPN1
R:371->G,S:372->G
VRSRVV
RC3H1
L:754->P
SLDELH
RRP7A
W:203->R
DEEGWV
SLC7A6OS
Y:253->C
YPEEES
SETD1A
L:496->P
IEMLLK
STX5
T:61->A,Q:62->R
RTQEFL
TAF6
V:463->A
LLCSQV
VPS13D
L:2900->P,W:2901->R
TGCTLW
WDR33
W:508->R
INTRON AWM
ZNF830
K:368->G
VKGALL
b
N=2622
Edited > 0.05: 62
EIF3G W12R
RTF1 F594P
WDR59 Y618H
WDR59 Y619H
ANKRD1 I2427V
Genomic Edit Frequency
Sensor Edit Frequency
0
1
0
1

**Supplementary figure 6: rhAmpSeq screen validates genomic locus editing**

**a**, Comparison of the *predicted, sensor* and *genomic* edits for 19 SLiM targets from the rhAmpSeq screen. Logos show the protein level edit frequency for the *sensor* (as a proportion of all edited and unedited sensor) and *genomic* (as a proportion of edited genomic locus) data. Low, indicates that the sample has low sequencing counts. Motifs in ANO8 and SOX12 had too low sequencing counts for quantification and are not shown. **b**, Comparison of observed sensor and genomic edit frequencies for all potential edits within the gRNA-targeted region across 23 targets in the rhAmpSeq screen. Each data point represents a specific potential protein-level mutation in the gRNA-targeted region. Predicted edits for gRNAs shown in Figure 3i are indicated.

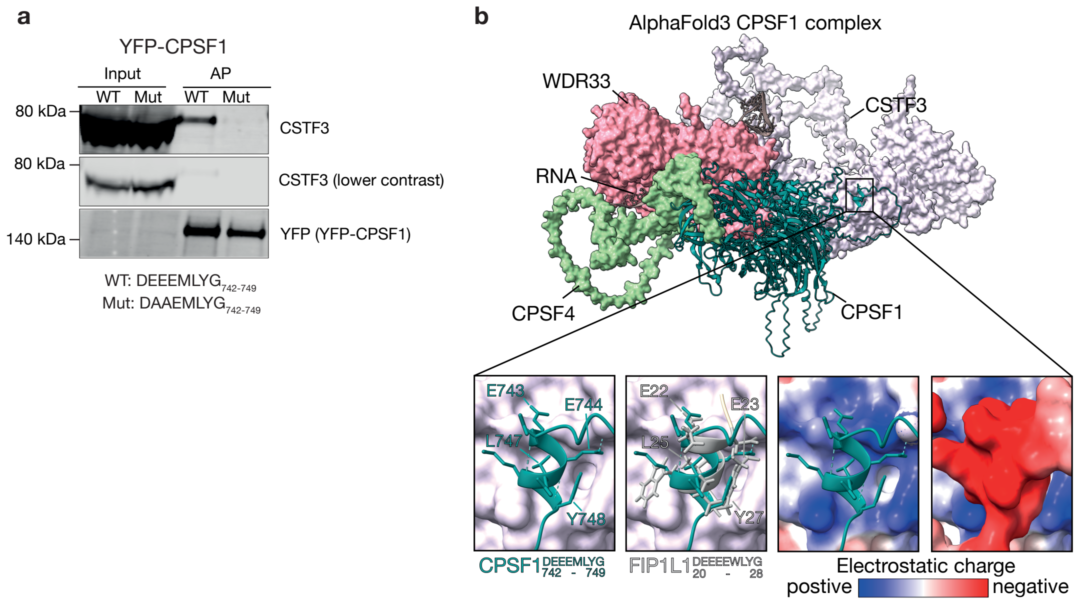


**Supplementary Figure 7: A novel SLiM in CSTF-CPSF cleavage and polyadenylation complex is required for complex formation.**

**a**, Western blot analysis of affinity purified (AP) full length YFP-CPSF1 wild-type (WT) and E743A/E744A mutant (Mut) from HeLa cells probed for CSTF3 to validate AP-MS result ( n=1). **b,** Top, AlphaFold3 prediction of WDR33-CSTF3-CPSF4-RNA-CPSF1 (PTM = 0.67, iPTM = 0.74), based on the pre-mRNA 3'-end processing machinery complex structure (PDB: 6URO). Bottom, details of prediction. Left square, detail of $CPSF1_{742-749}$ CSTF3 contact site. Middle left square, $CPSF1_{742-749}$ CSTF3 contact showing $FIP1L1_{20-28}$, aligned using the FIP1L1-CSTF3 crystal structure from PDB: 7ZY4. Middle right square, cartoon representation of $CPSF1_{742-749}$ with electrostatic surface representation of CSTF3. Right square, electrostatic surface representation of $CPSF1_{742-749}$ and CSTF3.

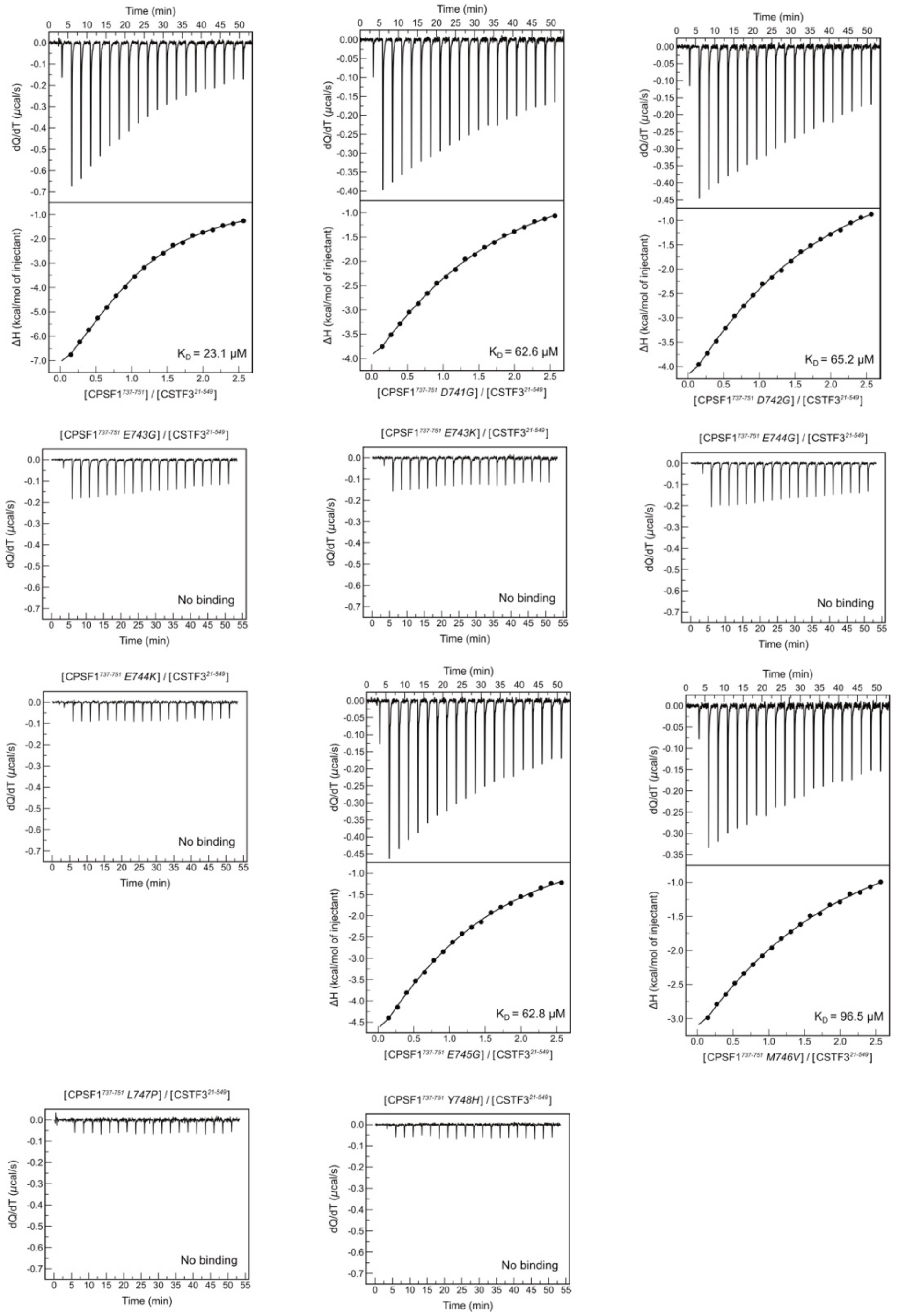


**Supplementary figure 8: Mutations in CPSF1 LYG motif cause loss of binding to CSTF3.** Isothermal calorimetry data for CPSF1$^{737\text{-}751}$ wild-type and single amino acid mutant peptides titrated against CSTF3$^{\textit{21-549}}$.

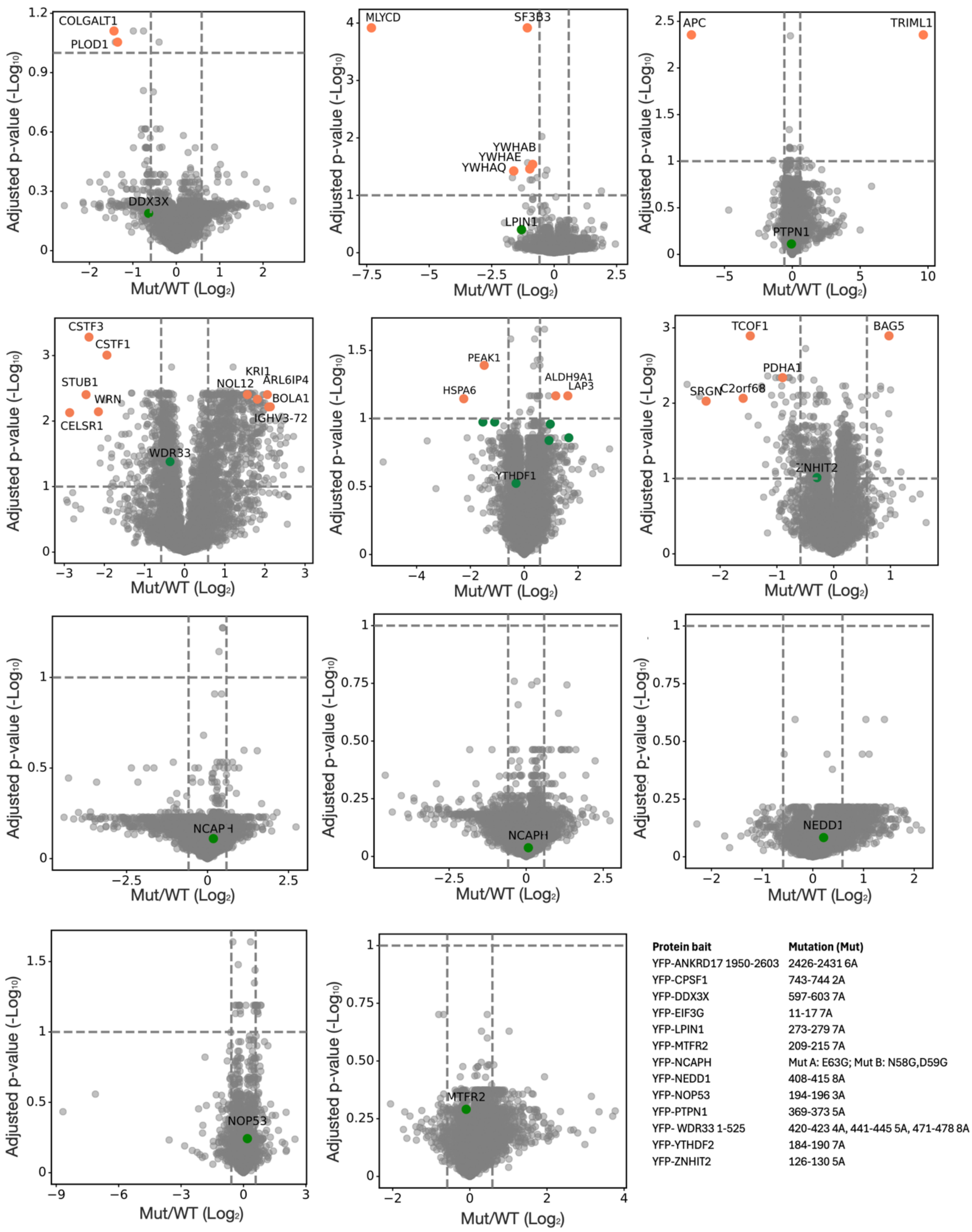


| Protein bait | Mutation (Mut) |
|---|---|
| YFP-ANKRD17 1950-2603 | 2426-2431 6A |
| YFP-CPSF1 | 743-744 2A |
| YFP-DDX3X | 597-603 7A |
| YFP-EIF3G | 11-17 7A |
| YFP-LPIN1 | 273-279 7A |
| YFP-MTFR2 | 209-215 7A |
| YFP-NCAPH | Mut A: E63G; Mut B: N58G,D59G |
| YFP-NEDD1 | 408-415 8A |
| YFP-NOP53 | 194-196 3A |
| YFP-PTPN1 | 369-373 5A |
| YFP- WDR33 1-525 | 420-423 4A, 441-445 5A, 471-478 8A |
| YFP-YTHDF2 | 184-190 7A |
| YFP-ZNHIT2 | 126-130 5A |

**Supplementary Figure 9: Mutation of predicted SLiMs disrupt protein interactions.**

AP-MS interactome data for YFP-tagged full length (or a large fragment) protein baits in a wild-type (WT) vs indicated SLiM mutant (Mut) version affinity purified from HeLa cells (n=4) . Adjusted *p*-value cut-offs are set to 0.1. Significantly lost or gained interactors are labeled orange (maximum 5). Baits are labeled green. Data for ANKRD17, CPSF1 and EIF3G are shown in main figures.

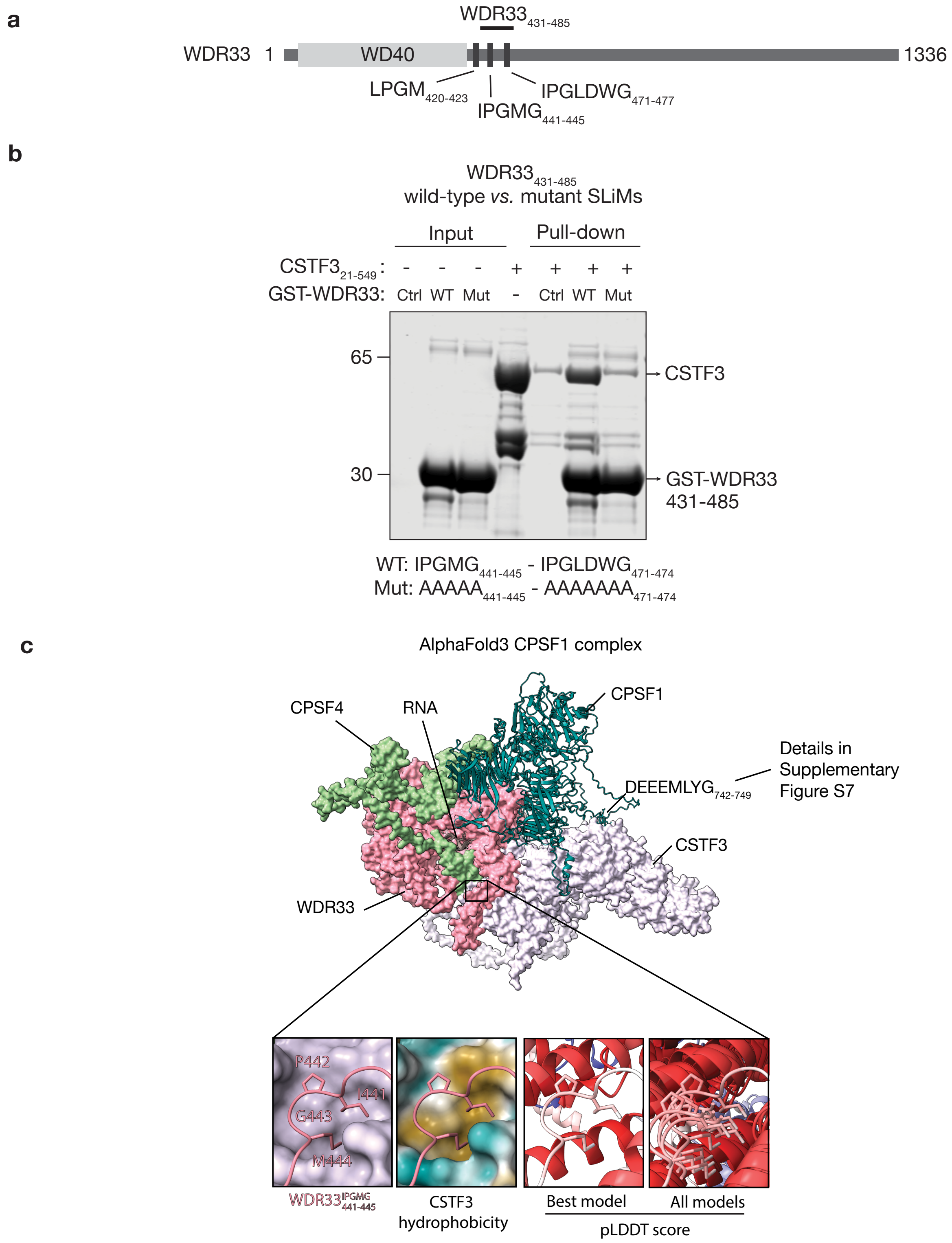
a
$WDR33_{431-485}$
WDR33 1
WD40
1336
$LPGM_{420-423}$
$IPGMG_{441-445}$
$IPGLDWG_{471-477}$
b
$WDR33_{431-485}$
wild-type *vs.* mutant SLiMs
Input
Pull-down
$CSTF3_{21-549}$: - - - + + + +
GST-WDR33: Ctrl WT Mut - Ctrl WT Mut
65
30
CSTF3
GST-WDR33
431-485
WT: $IPGMG_{441-445}$ - $IPGLDWG_{471-474}$
Mut: $AAAAA_{441-445}$ - $AAAAAAA_{471-474}$
c
AlphaFold3 CPSF1 complex
CPSF4
RNA
CPSF1
$DEEEMLYG_{742-749}$
Details in
Supplementary
Figure S7
CSTF3
WDR33
P442
I441
G443
M444
$WDR33^{IPGMG}_{441-445}$
CSTF3
hydrophobicity
Best model
All models
pLDDT score

**Supplementary Figure 10. SLiMs in WDR33 directly bind CSTF3**

**a,** Domain structure of WDR33 showing the two IPG (441-445 and 471-478) and one $LPGM_{420-423}$ SLiMs. **b**, GST pull down of wild-type (WT) or SLiM alanine mutant (Mut) GST-$WDR33_{431-485}$ with $CSTF3_{21-549}$, analysed by SDS-PAGE and InstantBlue staining (n=3). **c**, Details of WDR33 $IPGMG_{441-445}$ CSTF3 interaction observed in the AlphaFold3 prediction of WDR33-CSTF3-CPSF4-RNA-CPSF1 (PTM = 0.67, iPTM = 0.74) (tilted 90 degrees backwards as compared to Supplementary Figure S7b). Left square: WDR33 $IPGMG_{441-445}$ motif contact site with CSTF3. Middle left square: cartoon representation of $CPSF1_{742-749}$ with hydrophobic(orange)/hydrophilic(cyan) surface representation of CSTF3. Middle right square: best model pLDDT scores of WDR33 and CSTF3 represented as cartoon. Red indicates high pLDDT, blue indicates low pLDDT. Right square: Alignments of all models, represented as cartoon, showing pLDDT scores of WDR33 and CSTF3. For the best model, the complete sequence of WDR33 is shown. For clarity, the other models show CSTF3 in full but for WDR33 only $IPGMG_{441-445}$ is shown. Red indicates high pLDDT, blue indicates low pLDDT.

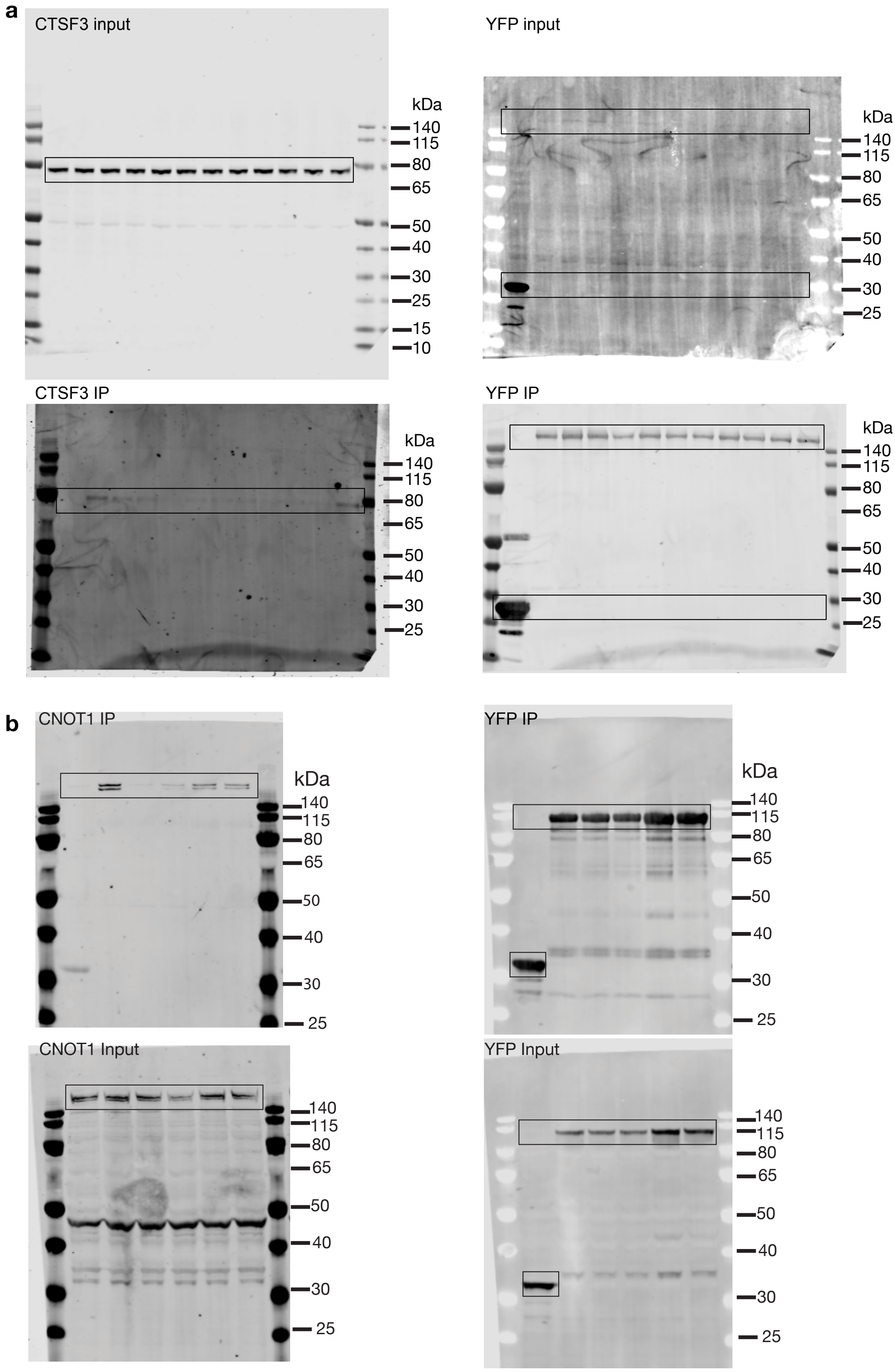


**Supplementary Figure 11: Uncropped Western blots**

**a,** Uncropped Western blots relating to Fig. 4e. **b,** Uncropped Western blots relating to Fig. 5i.
